# Transcription-induced mutation and ancient gBGC shape the evolution of human transcriptional start sites

**DOI:** 10.1101/2025.11.24.690122

**Authors:** Yi Qiu, Emilie Hebraud, Fanny Pouyet, Alexander F. Palazzo

**Affiliations:** Department of Biochemistry, University of Toronto, Toronto, Ontario, M5G1M1, Canada; Laboratoire Interdisciplinaire des Sciences du Numérique, Université Paris-Saclay, Gif-sur-Yvette, 91190, France; Institut Agro, Rennes Angers

**Keywords:** RNA Polymerase II, mutation bias, gBGC, transcriptional start site, CpG, evolution modeling

## Abstract

In the human genome, mutation rates vary along genes, yet how their fine-scale structure contributes to gene evolution has remained largely unexplored. Here, we map inherited mutations at single-nucleotide resolution and uncover a striking hypermutation peak at transcription start sites (TSSs). This pattern is observed both in population polymorphisms and in *de novo* mutations from parent-offspring trios, and is dependent on transcriptional activity in testes. We also find similar hypermutation peaks at TSSs used to produce long noncoding RNAs and at intergenic RNA Polymerase II pause sites. In addition, we identify distinct mutational signatures at exon-intron boundaries and in introns. By comparing the current nucleotide content to predicted equilibrium levels, inferred from mutation and fixation rates, and by analyzing derived ancestral frequency spectrums, we detect signals that are compatible with a low level of ongoing background GC-biased gene conversion (gBGC) throughout the region and ancestral elevated gBGC activity downstream from the TSS. Using a forward-in-time simulation algorithm, we show that the current nucleotide composition surrounding the TSS of protein-coding genes is best explained by local mutation biases coupled to ongoing and ancestral patterns of gene conversions. Our simulations allow us to infer several features of these gBGC events. Overall, these findings indicate that the nucleotide composition around TSSs is largely shaped by non-adaptive forces, particularly mutation bias and gBGC.

## BACKGROUND

One striking feature of transcriptional start sites (TSSs) of human protein-coding genes is that they have high GC-content (Zhang et al., 2004; Louie et al., 2003; Polak and Arndt, 2008; Palazzo and Kang, 2021). It is generally assumed that such major nucleotide features of the genome are solely the product of natural selection. Despite this, new findings have indicated that non-adaptive evolutionary forces may play a significant role in shaping nucleotide content, even at protein-coding genes. For example, ancient patterns of recombination around the TSS of human protein-coding genes likely caused these regions to be enriched in GC-content (Qiu et al., 2024) through the action of a strong non-adaptive force called GC-biased gene conversion (gBGC) (Duret and Galtier, 2009). gBGC happens at meiotic recombination initiation sites where maternal and paternal strands form mixed heteroduplexes. In these regions any mismatches due to heterozygosity are resolved in favor of G/C over A/T. This causes a bias for G/C single nucleotide polymorphisms (SNPs) to be passed down to the next generation over A/T SNPs (Duret and Galtier, 2009). Note that these recombination initiation events can be resolved to form either crossovers, or more often non-crossovers. Both processes likely promote gBGC. Although gBGC is largely directed away from the TSS in humans and rodents, most mammals have some degree of gBGC at the TSS (Joseph et al., 2024). Recombination tends to be concentrated at recombination hotspots. In most vertebrates, these hotspots are determined by PRDM9 and in its absence, recombination occurs preferentially at TSSs (Brick et al., 2012; Auton et al., 2013). Despite this, certain alleles of PRDM9 allow for some TSS-associated recombination (Hoge et al., 2024). Because the PRDM9 gene is rapidly evolving (Lesecque et al., 2014), and its recognition motifs are constantly destroyed by recombination, which is mutagenic (Hinch et al., 2023), TSSs likely experience dynamic cycles of GC-content change. When some TSS-associated recombination is allowed, due either to the presence of certain PRDM9 alleles, or an overall lack of PRDM9 motifs due to mutational decay, these regions rapidly gain GCs. When recombination is effectively directed away from TSSs, these regions slowly lose GCs.

Another factor that may affect the nucleotide content of TSSs is biased mutations, which can shape both adaptive and non-adaptive evolution (Stoltzfus and Yampolsky, 2009; Stoltzfus, 2021). TSSs are particularly mutation-prone because RNA Polymerase II accumulates at promoter-proximal pause sites (Jonkers and Lis, 2015; Core and Adelman, 2019), creating multiple sources of DNA damage (Gómez-González and Aguilera, 2019). Paused RNA Polymerase II causes local DNA unwinding and downstream supercoiling that topoisomerases must relieve by cleaving and re-ligating DNA. Failures in re-ligation generate double-stranded breaks and subsequent mutations during repair (Singh et al., 2020). Paused RNA Polymerase II also creates DNA replication fork roadblocks; collisions between paused polymerase and DNA replication machinery produce DNA damage events, including double-stranded breaks (Gómez-González and Aguilera, 2019). RNA Polymerase II pausing also promotes R-loop formation that elevates local mutation rates through multiple pathways (Gómez-González and Aguilera, 2019). R-loops sequester the template strand in DNA-RNA hybrids, which acts as roadblocks for DNA replication forks. At the same time, R-loops leave the coding strand single-stranded and exposed for extended periods, making it vulnerable to chemical damage (Sollier and Cimprich, 2015). All of these mechanisms increase the mutation rate exactly where GC-content is highest.

These mechanisms operating at the TSS may be compounded by broader mutational processes affecting gene bodies. The DNA damage generated by either topoisomerase activity, replication-transcription conflicts, or R-loops, promotes the activity of translesion DNA polymerases, which are error-prone (Goodman and Woodgate, 2013). Whether they are attempting repairs, or simply bypassing DNA lesions, these enzymes will introduce secondary mutations that extend far away from the original damage site, potentially affecting gene bodies. Certain repair enzymes are recruited by histone modifications associated with exons, and this may lead to the accumulation of differential mutations between exons and introns (Huang et al., 2018). Moreover, transcription elongation rates are slower over exons than introns (Jonkers and Lis, 2015) and this may further allow mutation patterns to differ in these two regions. There are also transcription-coupled DNA repair pathways that identify and repair lesions found solely on the template strand (Fousteri and Mullenders, 2008). As a result, certain mutations occur more often on the coding strand and thus create strand-asymmetric mutational patterns across gene bodies.

Although most of these investigations have been conducted in somatic cells (both normal and cancerous), large scale studies indicate that transcription and DNA replication have major impacts on the rate of mutations in germ cells, thus impacting evolution (Seplyarskiy and Sunyaev, 2021; Seplyarskiy et al., 2021; Cortés Guzmán et al., 2025). Despite all of this, the contribution of mutational bias in shaping genomic content remains underappreciated by most molecular biologists (Palazzo and Kejiou, 2022).

Here we examine mutation rates with single nucleotide resolution, revealing a hypermutational signal at the TSS of protein-coding genes. We show that these peaks correlate with RNA Polymerase II-associated transcription in germ cells. Comparing equilibrium nucleotide content calculated based on human mutation rates and substitution rates, and determining how mutations have recently spread by analyzing derived ancestral frequencies, we estimate the contribution of mutational bias and gBGC in shaping nucleotide content at the TSS in humans. We then validate the effects of local mutation and fluctuations in gBGC on the nucleotide content of protein-coding genes through a simulation of gene evolution.

## RESULTS

### The TSSs of human protein-coding genes experience elevated rates of mutations

To determine whether trends in nucleotide content around the transcriptional start site (TSS) of protein-coding genes (current average nucleotide content is shown in Figure 1A) are shaped in part by localized mutational hotspots, we identified low frequency single nucleotide polymorphisms (SNPs) in the human population using the gnomAD database (Karczewski et al., 2020). Low frequency SNPs are generally evolutionarily young, having been created by recent mutational events, and hence have not experienced significant selection pressure, and thus can be used to infer the base mutation rate. The gnomAD database contains half a billion low frequency SNPs (frequency <1%), isolated from 71,702 individuals, representing one SNP for every six nucleotides. The high density of data allowed us to infer the mutation rate at single nucleotide resolution around the TSS, which we mapped using Cap Analysis Gene Expression (CAGE) sequencing. Since only germline mutations are transmitted across generations and are thus the most relevant for evolution, we used CAGE sequencing from human testes (Noguchi et al., 2017). To infer local mutation rates, we defined the parameter *M*, based on the density of rare variants. The number of these low frequency variants depends not only on the mutation rate (*µ*) but also on population genetics factors such as the effective population size (*N_e_*), the selection coefficient (*s*) and the conversion bias due to gBGC (*b*) which affect not only the probability to observe these mutations but also their residence time in the 1% frequency bin. We therefore rescaled *M* so that its value in intergenic regions corresponds to the mutation rate, *µ*, inferred by *de novo* mutations as determined by parent-offspring trio sequencing (Palsson et al., 2025). We expect that the joint effects of *N_e_*, *b*, and *s* are minimal for very low frequency SNPs, thus making localized values of *M* ≈ *µ*. Under the assumptions of neutrality and constant population size, the rescaling factor (calculated to be 120 using intergenic regions, see Methods under “Rare SNP mapping and derivation of *M*”) can be interpreted as the approximate average number of generations ago these SNPs appeared in the human population.

**Figure 1.**
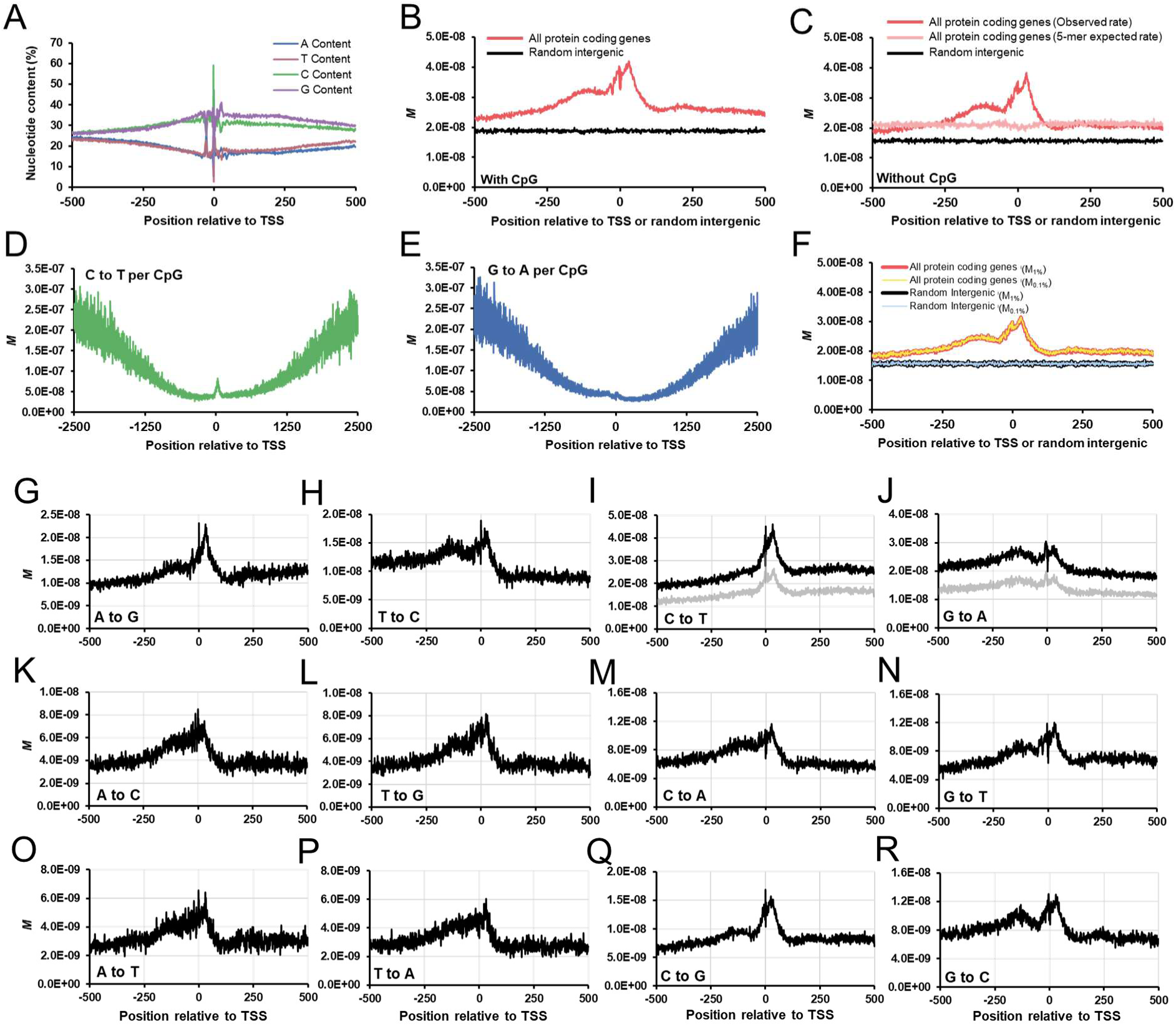
Single nucleotide resolution *M* for regions surrounding human protein-coding gene TSSs. A) Nucleotide content around the TSS, averaged over all human protein-coding genes and displayed at single nucleotide resolution. The TSS is positioned at 0. B) Single nucleotide resolution of *M_1%_*, over all TSSs from human protein-coding genes (red) or an equivalent number of intergenic regions (black) using low frequency SNP analysis from gnomAD. C) Single nucleotide resolution of *M_1%_* for nonCpG mutations, over all TSSs from human protein-coding genes (red) or an equivalent number of intergenic regions (black) using low frequency SNP analysis from gnomAD. Single nucleotide predicted rate based on 5-mer sequence context over all TSSs from human protein-coding genes (pink). D-E) Single nucleotide resolution of *M_1%_*, for CpG to TpG (D) and CpG to CpA (E) around the TSS of all human protein-coding genes. F) Comparison of single nucleotide resolution of *M_1%_* (red for TSS regions, black for intergenic regions) and *M_0.1%_* (yellow for TSS regions, blue for intergenic regions). G-R) Single nucleotide resolution of *M_1%_*, for each indicated mutation type around the TSS of all human protein-coding genes. For I-J total *M_1%_* is shown in black, nonCpG *M_1%_* is shown in grey.

Using *M,* we estimated the mutation rate surrounding the TSS of human protein-coding genes with (Figure 1B), and without CpGs (Figure 1C). Strikingly, we observed a peak in *M* in the region surrounding the TSS. In agreement with the idea that transcription is mutagenic, we observed that *M* around the TSS was higher than in intergenic regions. When we excluded CpGs, *M* was above the expected context rate (Figure 1C) as calculated using 5-mer counts and the rate at which each pentamer acquires low frequency SNPs in intergenic regions (see Methods under “5-mer sequence context rate calculation”). Note that the expected context rate was much higher when CpGs were included (Supplemental Figure 1B) and this is to be expected as CpGs near the TSS are hypomethylated and protected from mutational decay (as seen in Figure 1D and 1E). The suppression in CpG mutation is in agreement with previous substitution analyses (Polak and Arndt, 2008). Note that non-methylated C deaminates to U and is much more readily repaired than 5meC, which deaminates to T (Sved and Bird, 1990; Walsh and Xu, 2006). The lack of methylation around the TSS allows deaminated CpGs in these regions to be recognized and efficiently reverted back to their original state.

The increase in *M* at the TSS consisted of two distinct peaks. The major peak was centered 25 nucleotides downstream of the TSS, while a minor one was centered at about 110 nucleotides upstream from the TSS. Furthermore, we detected a relative depression in *M* centered around 140 nucleotides downstream of the TSS followed by another shallow peak centered at 200.

To determine whether *b* or *s* was significantly altering *M*, we recalculated this metric using ultralow frequency SNPs (<0.1%), which have on average been in the population for fewer generations and experienced less selection or gBGC. The two curves (*M_1%_* and *M_0.1%_*) were superimposable for both TSS and intergenic regions (Figure 1F), indicating that the effects of *b* and *s* are indeed minimal.

Next, we examined the effects of *M* on GC-content (note that by default all *M* in this study is *M_1%_* unless specified). In agreement with our previous analysis of *de novo* mutations (Qiu et al., 2024), which were inferred from parent-offspring whole genome sequencing (Jónsson et al., 2017), we find a net loss of Gs and Cs around the TSS (Supplemental Figure 1A). Within this greater trend, there are two exceptions. First, there are fewer GC-losses in a region 23 nucleotides upstream of the TSS, a location that corresponds to the TATA box, which is relatively GC-poor. Second, there are fewer GC-losses right at the TSS and this may be due to the particular distribution of nucleotides at the +1 and –1 positions relative to the TSS (Carninci et al., 2005).

Next, we analyzed *M* for different mutation types (Figure 1G-R) including non-CpG mutations to TpG and CpA (Figure 1I-J grey line). The major peak (25 nucleotides downstream from the TSS) is seen in all mutation types. Interestingly, the upstream minor peak (at ∼110 nucleotides upstream from the TSS) is only seen in a subset of mutation types. For example, it is not readily detectable in C to T transition mutations (Figure 1I), but is prominent with all mutations away from G (i.e., G to A, G to T and G to C, Figure 1J, N and R). We could also see a small peak near the TSS in the CpG *M* analysis (see small peaks near “0” in Figures 1D-E) that mirrored the peak in total and non-CpG *M* graphs (Supplemental Figure 1C-D), suggesting that these are all a product of the same process.

Looking more broadly, our measurements of *M* indicate that regions downstream of the TSS (>200 nucleotides), which are mostly intronic regions, have mutation patterns that differ from upstream regions (>200 nucleotides upstream from the TSS). For example, downstream regions have lower T to C mutations than upstream regions (Figure 1H). Notably, these patterns are not strand symmetric (i.e. regional changes in the T to C mutation rate are not mirrored in the A to G mutation rate – compare Figures 1H to 1G). In the absence of selection, this is expected to produce strand asymmetry, also known as skew. This is consistent with the idea that the nucleotide distributions in human introns are due to subtle differences in the nucleotide mutation or repair rates between the template and coding strands (Touchon et al., 2004; Polak and Arndt, 2008; Seplyarskiy and Sunyaev, 2021; Palazzo et al., 2024).

We also analyzed somatic mutations around the TSS of protein coding genes based on the SomaMutDB (Supplemental Figure 1E-F). We excluded datasets from samples with DNA damage repair defects since these would cause additional biases. In this dataset, we also observe a hypermutational peak from somatic mutations around the TSS. However, we only observe one prominent peak centred ∼30 nucleotides downstream of the TSS, mirroring the main peak in *M* derived from germline mutations (Figure 1B-C).

Using 50 base pair windows spanning the 1kb region surrounding the TSS of protein coding genes or random intergenic sequences, we assessed SBS mutational signatures based on rare SNPs (<1%) (Suppleental Figure 2) according to COSMIC-SigProfiler (Tate et al., 2019; Díaz-Gay et al., 2023). We performed this analysis either normalizing by the trinucleotide frequencies of each region (Supplemental Figure 2A), or without normalization (Supplemental Figure 2B). Normalization altered the percentage distribution of SBS signatures of each window but mostly retained the observed SBS types. While intergenic regions had mostly SBS5, consistent with previous analyses of germline mutations (Rahbari et al., 2016), TSS of protein-coding genes had additional signatures, indicating an enrichment of mutational processes associated with transcription-associated damage and DNA repair.

**Figure 2.**
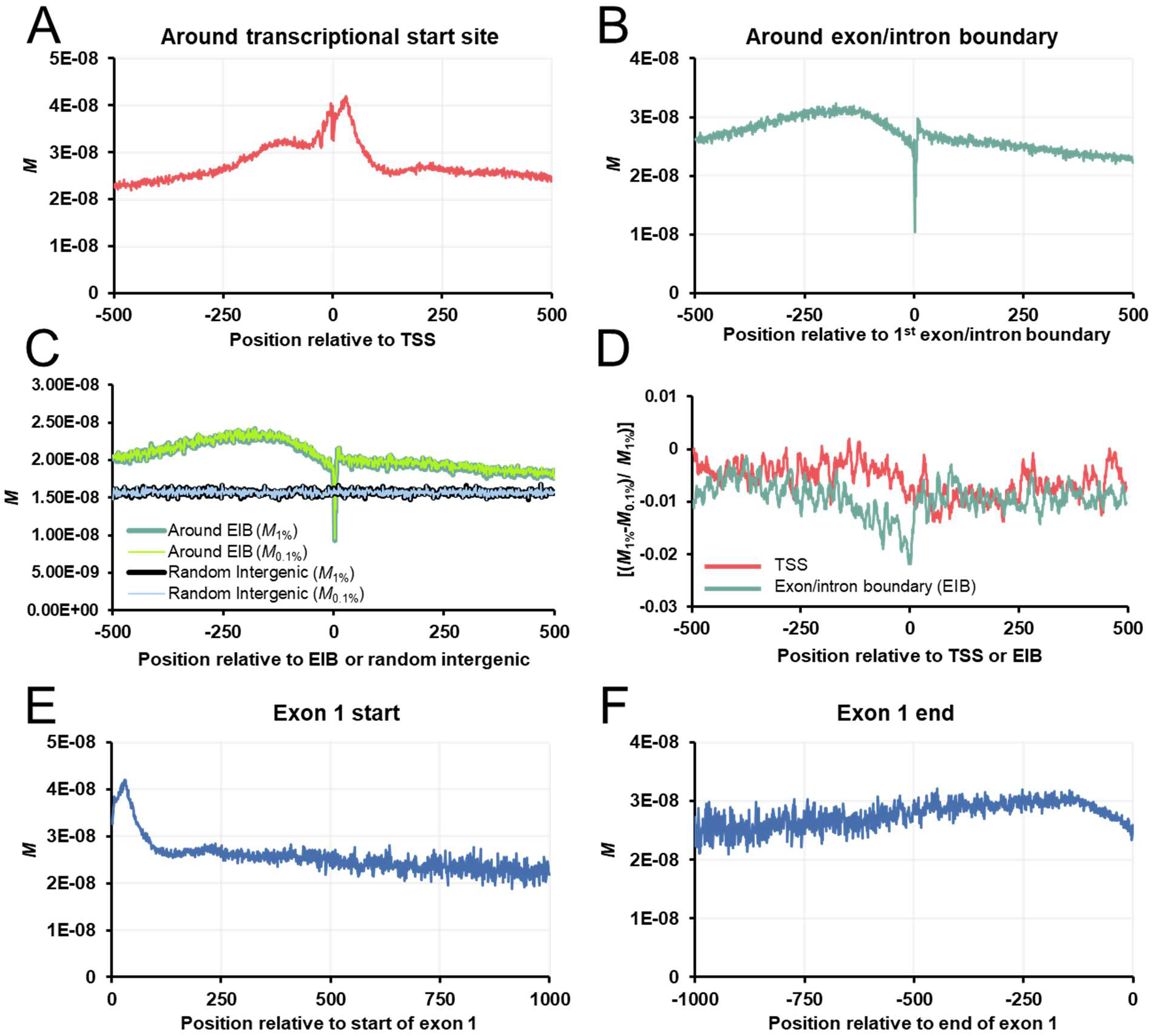
Single nucleotide resolution *M* for the first exon of human protein-coding genes. A-B) Single nucleotide resolution of *M_1%_*, over all TSSs (A), or the first exon/intron boundary (B), of all human protein-coding genes. C) Comparison of single nucleotide resolution of *M_1%_* (dark green for exon/intron boundaries or “EIB”, black for intergenic regions) and *M_0.1%_* (light green for exon/intron boundaries, blue for intergenic regions). D) Fractional change between *M_1%_* and *M_0.1%_* for regions surrounding the TSS and EIB. Negative numbers indicate a depletion of SNPs between 0.1% and 1% as compared to the expected value. Note that CpGs were omitted (see methods). Also note that the greatest change occurs right at the exon/intron boundary where 5’ splice site motifs are located. E) Single nucleotide resolution of *M_1%_* over all first exons aligned so that the first nucleotides are positioned at 0. F) Single nucleotide resolution of *M_1%_* over all first exons aligned so that the last nucleotides (at the EIB) are positioned at 0.

We observed an enhancement of SBS23, which has unknown etiology but is enriched in C to T mutations. This signature is similar to SBS11, SBS19, and SBS30 (see Supplemental Figure 2C), which were previously reported around TSSs in the germline (Cortés Guzmán et al., 2025), although not directly detected in our analysis. We also observe SBS46, which is similar to SBS12 (enriched in T to C mutations) (Supplemental Figure 2C), that was associated with replication-strand asymmetry, and also detected for germline mutations around TSSs by others (Cortés Guzmán et al., 2025).

Furthermore, we observe SBS3, SBS39 and SBS6. These are linked to distinct DNA repair pathways, including defective homologous recombination DNA damage repair (SBS3), double-stranded DNA breaks repair through non-homologous end joining (NHEJ) or microhomology-mediated end joining (SBS39) (Ding et al., 2024), and defective DNA mismatch repair (SBS6) (Németh et al., 2020). SBS3 was previously reported near TSSs for germline mutations (Cortés Guzmán et al., 2025). SBS39, which is enriched in C to G mutations, has previously been associated with homologous recombination deficiencies (Degasperi et al., 2020), was also found in Cortés Guzmán et al. (Cortés Guzmán et al., 2025) and resembles germline component 8 detected in a separate study (Seplyarskiy et al., 2021). We also observe SBS98, which is enriched in CpG to NpG mutations, and resembles germline mutation signature component 11 in the same study (Seplyarskiy et al., 2021).

Together, these results support the idea that TSS regions are enriched for multiple mutational processes linked to replication asymmetry and DNA damage repair.

### The main mutational hotspot downstream of the TSS occurs within the first exon

To determine where the main mutational hotspot lies, with respect to other gene features, we reanalyzed *M* for all mutations (Figure 2A) and the twelve individual mutation types (see Supplemental Figure 3), with respect to exon and intron boundaries.

When protein-coding genes were lined up so that the boundary between their first exons and first introns were aligned (Figure 2B), we could see a dip in *M* right at the boundary. This is likely due to the fact that mutations in many 5’ Splice Sites (5’SS), which recruit the spliceosome to the 5’ end of the intron, are under high levels of selection (Christmas et al., 2023), and thus under-represented in the rare SNP pool. Upstream of the exon/intron boundary, *M* increases – this is likely due to the spreading out of the main peak in *M* that we previously mapped 25 nucleotides downstream of the TSS (see Figure 2A). Right upstream of the exon/intron boundary (relative position around –125 to 0), we see a very slight decrease in *M* (Figure 2B). It is possible that this slight decrease is due to the elimination of highly deleterious non-synonymous mutations in coding regions from gnomAD. Within the intron we observed a steady decrease in *M* although this seems to be true for some classes of mutations (for example C to A, Supplemental Figure 3G column 4) but not others (for example A to G, Supplemental Figure 3A column 4).

To gage how selection is affecting these regions we recalculated *M* for ultrarare SNPs (<0.1% frequency) to look for changes. Again, the two graphs were almost completely superimposable (Figure 2C). When we measured the fractional difference between *M_1%_* and *M_0.1%_*, the biggest dip (amounting to a change in 2%) was right at the exon-intron boundary where the 5’SS is found (Figure 2D). This change was larger than at any other region, including around the TSS (Figure 2D) and is in agreement with the idea that 5’SSs are under negative selective pressure. Other differences could be seen upstream of the exon-intron boundary indicating a lower level of selection in these regions which encompass coding regions in the first exons.

Next, we lined up all first exons while excluding all intronic sequences and recalculated *M* (“Exon 1 start”). We observed a clear signal at 25 nucleotides downstream from the TSS (Figure 2E). This could also be seen for all mutation types (Supplemental Figure 3A-L column 2), suggesting that the main peak in *M* was not due to the elevation of a particular type of mutation. At longer distances away from the TSS, we see a drop in *M* that is similar to what is seen in introns (compare the slope in Figure 2E, of nucleotides >250 downstream of the exon 1 start, to the slope of Figure 2B, of nucleotides downstream of the exon/intron boundary). When exons were lined up so that the last nucleotides of exon 1 are aligned (“Exon 1 end”, all alignments start at 0 on the right side of the *x-axis* in Figure 2F), we saw a decline in *M* that starts at 150 nucleotides upstream of the exon/intron boundary and continues to fall as one approaches the boundary (Figure 2F). Again, this is true for all mutation types (Supplemental Figure 3, column 3).

From this analysis, we concluded that the overall patterns of *M* in the exons are more complicated than those seen in the introns. Some patterns are mutation type specific, others are found across all mutation types.

### Mutational spectra associated with protein-coding TSSs are absent in genes not expressed in the testes

The association of distinct patterns of *M* around the TSS supports the notion that the act of transcription may be mutagenic, as has been documented in normal and cancerous somatic cells (Gómez-González and Aguilera, 2019). If this was true, and since only germline mutations are transmitted, the patterns of *M* that we observe are not expected to be present in genes that are transcriptionally silent in the germline or in gametes. Moreover, since about 70% of all mutations occur in the paternal chromosomes (Jónsson et al., 2017), we hypothesized that these mutations would be more pronounced in DNA regions that are undergoing transcription in the testes. In agreement with our predictions, *M* compiled from genes not expressed in testes lacked the hypermutation peak (Figure 3A, compare red and blue lines, see statistical analysis for *M* at the TSS in Figure 3B, red and blue bars). This was true regardless of whether CpG were included (Figure 3A) or excluded (Supplemental Figure 4C). *M* compiled from genes not expressed in testes are similar to the predicted context rate, which was different than what we had seen in protein-coding genes (compare Supplemental Figure 4D to Supplemental Figure 4E).

**Figure 3.**
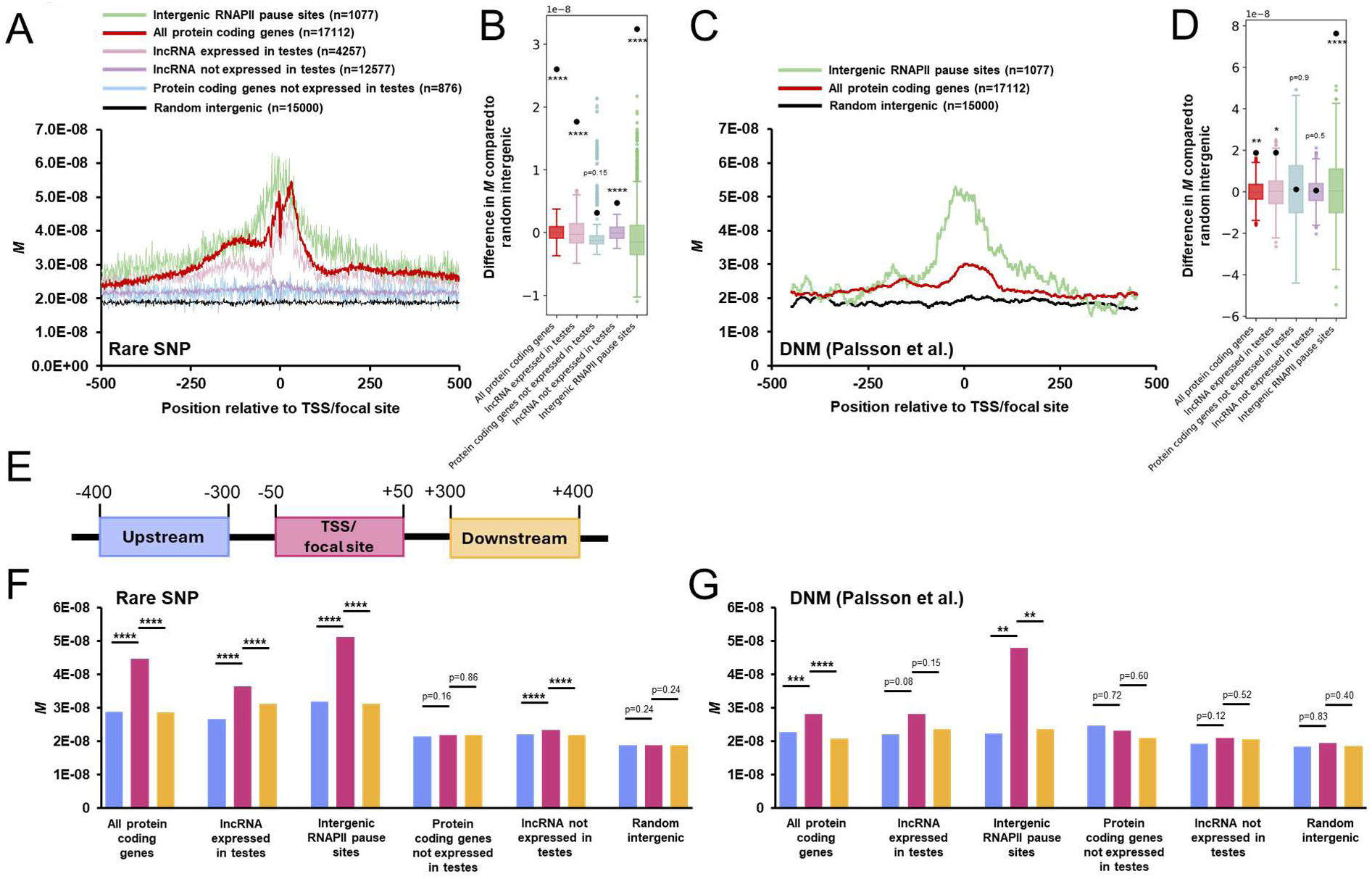
Hypermutation signals around human protein-coding gene TSSs correlate with transcription in the testes. A) Single nucleotide resolution of *M_1%_*, over various indicated regions from the human genome. B) Permutation test results comparing *M_1%_* for a 100 bp window around the 0 point for various indicated regions, against random intergenic regions. Distribution of differences from 1000 randomized permutations are displayed in box and whisker plots. Actual differences are plotted with black dots with *p* values indicated. C) Mutation rates, averaged over all indicated regions from the human genome, based on *de novo* mutations as inferred by parent-offspring trio sequence analysis. Rates are plotted as a sliding window of 100 nucleotides. D) Permutation test results comparing mutations rates as inferred by parent-offspring trio sequence analysis for a 100 bp window around the 0 point averaged over various indicated regions, against random intergenic regions. Distribution of differences from 1000 randomized permutations are displayed in box and whisker plots. Actual differences are plotted with black dots with *p* values indicated. E) Diagram depicting analyzed zones used in (F-G) to determine the presence of a mutational peak. F) *M_1%_* over 100 bp windows as indicated in (E) for various indicated regions. *p* values between the center and upstream or downstream regions are indicated. G) Mutation rates, averaged over 100 bp windows as indicated in (E) for various indicated regions. *p* values between the center and upstream or downstream regions are indicated. *: p < 0.05; **: p < 0.01; ***: p < 0.001; ****: p < 0.0001.

Additionally, using rare SNPs (<1%), we assessed the SBS mutational signatures associated with the TSS of genes not expressed in testes. Depending on how the analysis was performed, SBS5 was the predominant signature. Despite this, we observe some of the main signatures generally associated with TSSs protein-coding genes (Supplemental Figure 2A-B). This could be due to either a low level of transcription in testes (below our cutoff), transcription in post-zygotic cells, or the low number of mutations in each bin (due to the paucity of genes in the non-expressed in testes pool, see Figure 3A legend) that reduces the signal.

To validate this observation, we compiled *de novo* mutations from two previous studies (Palsson et al., 2025; Seplyarskiy et al., 2023) and determined the relative mutation rate (Supplemental Figure 4A, note that the two datasets had some overlapping data, see Supplemental Figure 4B). Again, we observed the two main peaks in the mutation rate surrounding the TSS of human protein-coding genes. To determine the real rate, we limited ourselves to the Palsson et al dataset, which was collected from the sequencing of 5,420 parent-offspring genome trios (Palsson et al., 2025), as we did not have the number of individuals used to generate the second data set. Again, we observed both peaks (Figure 3C). The main peak was significantly above the intergenic rate (Figure 3C black line, see statistical analysis in Figure 3D red bar).

We then examined genes not expressed in testes. Unfortunately, the low number of these genes (Figure 3A legend), combined with the paucity of mutations in parent-offspring genome sequencing (Supplemental Table 1), made this data very noisy. Despite this, there was no difference between mutation rates at the TSS of genes not expressed in testes compared to intergenic regions (see Figure 3D blue bar), although the variance in the data was large (see large standard deviation whiskers for this gene set, blue bar in Figure 3D).

To further validate this data, we calculated either *M* from rare SNPs or mutation rates from *de novo* mutations (from parent-offspring genome trio analysis) at the TSS or in upstream and downstream regions. *Bona fide* peaks should have elevated metrics at the TSS compared to nearby regions. To increase the amount of data we looked at 100 nucleotide windows (+/− 50 nucleotides from the TSS, –300 to –400 nucleotides for upstream sequences and 300 to 400 nucleotides for downstream sequences; Figure 3E). For both *M* (rare SNP, Figure 3F) and *de novo* mutations from parent-offspring sequencing (DNM, Figure 3G), protein-coding genes in general had a hypermutation peak signal associated with the TSS (Figure 3F-G). This peak was also seen when *M* was calculated for all individual mutation types (Supplemental Figure 5). In contrast, protein-coding genes not expressed in testes had no difference for overall mutations (Figure 3F-G) or for any specific mutation (Supplemental Figure 5).

From this data we conclude that genes not expressed in testes lack the mutational signal seen at the TSS of most protein-coding genes.

### Mutational spectra associated with protein-coding TSSs are also seen in regions expressing lncRNAs in testes

To further validate the idea that these trends in *M* are due to transcription-based mutagenesis, we examined TSSs associated with regions producing annotated lncRNAs in testes. Note that most lncRNAs are transcribed by RNA Polymerase II. Remarkably, these regions had the same overall trend in *M* as protein-coding genes (Figure 3A pink line, see statistics in Figure 3B pink bar, also Figure 3F). Again, this inferred hypermutational peak is above the predicted mutation rate based on 5-mer context (Supplemental Figure 4F). When the mutation rate around these TSSs was examined using *de novo* mutations from parent-offspring sequencing, we observed a significant difference when these were compared to intergenic regions (Figure 3D pink bar, 3G). Although the peak was present when compared to upstream and downstream sequences (Figure 3F-G), it was only significant in the *M* analysis and this was likely due to both a paucity of data and high variability between lncRNA TSS regions in the analysis of *de novo* mutations. When we examined *M* for individual mutation rates, we could detect the main hypermutation peak for each mutation type (Supplemental Figure 5). We then examined *M* around TSSs of regions transcribed into annotated lncRNAs that are not expressed in testes. The *M* in these regions was mostly flat (Figure 3A, purple line), although there was a minute peak that was statistically higher than in intergenic regions (Figure 3B purple bar, also Figure 3F). We suspect this minute peak might be due to low levels of germline expression still present in these regions that were not effectively filtered out. When we examined *de novo* mutations from parent-offspring trios there was no significant peak present (Figure 3D purple bar, Figure 3G). Interestingly, these regions had slightly lower *M* than what is predicted by the 5-mer context (Supplemental Figure 4G). Note that the same trend was seen in genes not expressed in testes (Supplemental Figure 4E).

Using rare SNPs (<1%), we calculated the SBS signatures 1kb around lncRNA TSSs. We observed many of the same signatures that we had previously seen in protein-coding genes (Supplemental Figure 2). For TSSs of lncRNAs that are not expressed in testes these signatures were somewhat supressed and replaced with SBS5, although the degree of suppression depended on the exact analysis used.

The many resemblances that we have documented between TSSs for regions transcribed into lncRNAs and protein-coding genes, indicates that both regions undergo the same mutational biases.

### Intergenic RNA Polymerase II pause sites are also associated with hypermutation

One of the main reasons that transcription is thought to be mutagenic is that RNA Polymerase II tends to pause just downstream of the TSS, which can cause mutagenesis by several mechanisms (Gómez-González and Aguilera, 2019).

With this in mind, we analyzed *M* surrounding intergenic RNA Polymerase II pause sites, as mapped in testes. Note that these pause sites were filtered to exclude any that started near a protein-coding gene or a region transcribed into lncRNA to avoid using alternative TSSs of *bona fide* genes. As such, these sites likely represent non-functional regions that evolved motifs that have biochemical activity through drift (Struhl, 2007; Palazzo and Lee, 2015; Palazzo and Kejiou, 2022; Gvozdenov et al., 2023). Unlike TSSs, we do not have single nucleotide resolution for the exact start site and we do not have strand-specificity, thus all sites were analyzed on the + strand and represent a mixture of sequence from template and coding strands (with respect to RNA Polymerase II). Remarkably, intergenic RNA Polymerase II pause sites have *M* values and de novo mutation rates that exceed what we measured at protein-coding or lncRNA TSSs (Figure 3A-G, Supplemental Figure 4A). The same inferred hypermutational signal is also observed at these sites when only non-CpG SNPs are analyzed (Supplemental Figure 4C). As with the other regions, *M* for RNA Polymerase II pause sites is higher than what is predicted based on 5-mer sequence context (Supplemental Figure 4H). Note that the pattern of *M* around intergenic RNA Polymerase II pause sites appears symmetrical (Figure 3A), which is expected given that we do not have strand information.

We next examined the SBS signatures 1kb around RNA Polymerase II pause sites. Again we observed the same signatures as seen in protein coding genes and lncRNAs, with enriched proportions of SBS39 (Supplemental Figure 2).

From this data, we conclude that the presence of paused RNA Polymerase II in the testes genome is sufficient to promote an increase in inherited mutations. These mutations appear to have similarities to those that appear around the TSS of protein-coding genes.

### Equilibrium nucleotide contents suggest ongoing background gBGC and ancestral TSS-associated hotspots of gBGC

We used *M*, to calculate the expected equilibrium level for all four nucleotides at each position in the vicinity of human protein-coding TSSs and compared these to the observed nucleotide levels. To simplify our calculations, we only examined non-CpG positions and non-CpG *M*. We also calculated the expected equilibrium nucleotide level based on substitution rates as calculated from comparing human and chimpanzee sequences, using gorilla as an outgroup. We plotted equilibrium levels from both human and chimpanzee. Although they were both noisy due to the paucity of data, they resembled each other. As expected from our previously published results (Qiu et al., 2024), which indicated that the GC-peak is under decay at a rate consistent with neutral evolution, the current levels of G and C in the vicinity of the TSS are significantly higher than the predicted equilibrium levels based on the substitution rate (Figure 4A-D, compare black and blue lines).

**Figure 4.**
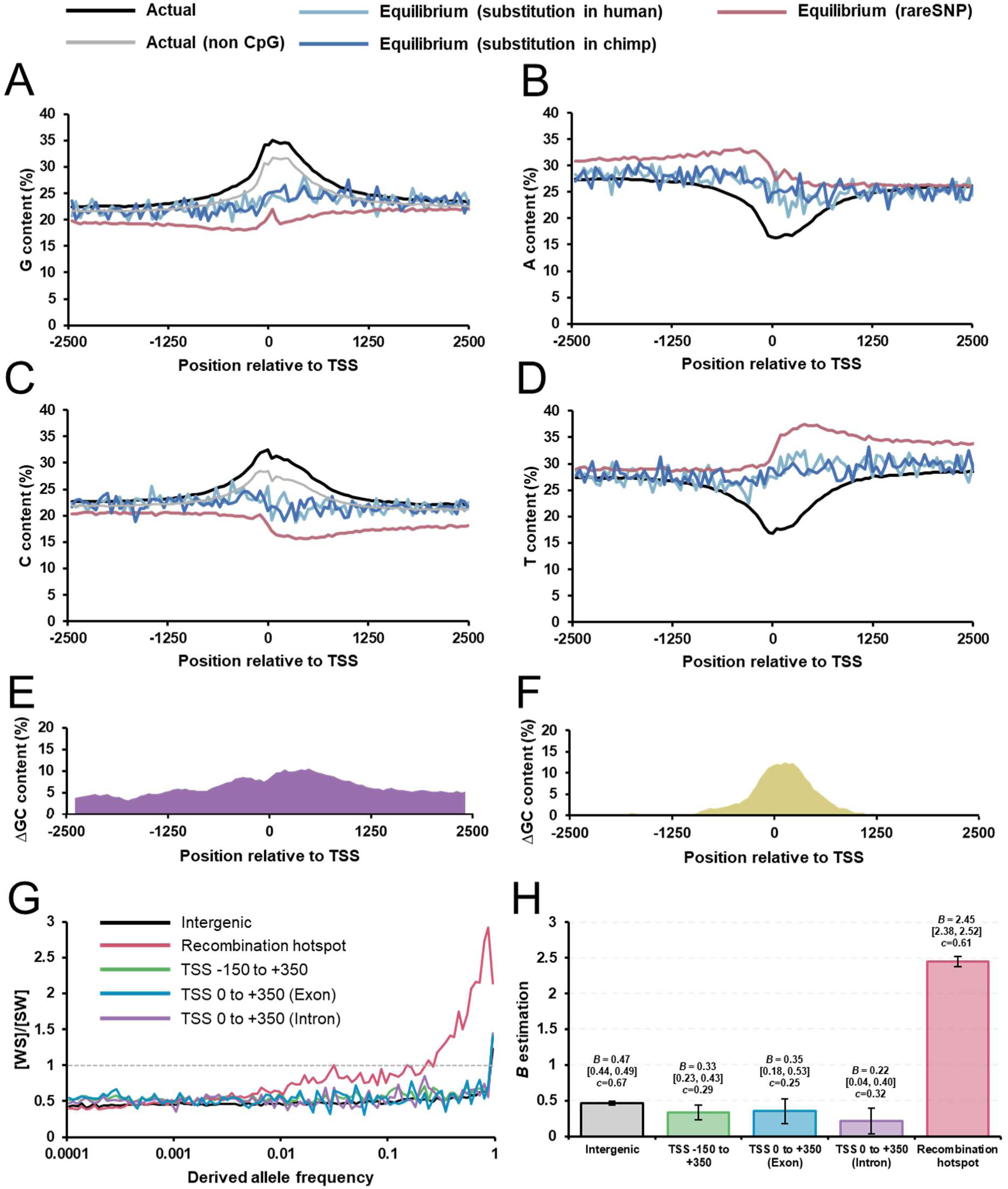
Equilibrium analysis and derived allele frequency spectra reveal ancestral, not ongoing, TSS-associated gBGC. A-D) The observed (black lines) nucleotide content and predicted equilibrium nucleotide levels based on either *M_1%_* (pink lines) or substitution rates inferred from human-chimpanzee-gorilla trio analysis (blue lines) for regions surrounding the TSS of human protein-coding genes. All plots use bins of 25 nucleotides. E) Difference in GC-content between predicted equilibrium levels based on substitution rates (averaged between human and chimpanzee) and *M_1%_* for regions surrounding the TSS of human protein-coding genes. The values use a sliding window of 125 base pairs. Note that the difference is uniform across the region, suggesting that background gBGC is acting throughout this region. F) Difference in GC-content between current nucleotide content in the human genome and predicted equilibrium levels based on substitution rates (averaged between human and chimpanzee rates) for regions surrounding the TSS of human protein-coding genes. The values use a sliding window of 125 base pairs. Note that the difference is most pronounced in a peak centered just downstream of the TSS of the graph, suggesting that TSS-associated gBGC occurred before the divergence of humans and chimpanzees. G) Ratio of weak-to-strong (WS) over strong-to-weak (SW) variants across derived allele frequency (DAF) bins for intergenic regions (black), recombination hotspots (pink), and the TSS −150 to +350 window partitioned into total, exonic, and intronic positions (green, blue, purple). Variants were polarized to ancestral and derived alleles using Ensembl’s primate ancestral sequence. H) Maximum-likelihood estimates of the population-scaled gBGC coefficient *B* for each region, with 95% confidence intervals from profile likelihood. *B* was fit jointly with a baseline mutation-bias parameter c that captures region-specific differences in mutation rate and base composition independent of DAF. Numerical values for *B* and c are shown above each bar.

To validate the results obtained with *M*, we repeated the equilibrium calculations using *de novo* mutations from the datasets we had previously analyzed (Seplyarskiy et al., 2023; Palsson et al., 2025). Even though these produced very noisy graphs (due to the paucity of data, Supplemental Table 1), they generally resembled the equilibriums calculated by *M* (Supplemental Figure 6).

Differences between mutational equilibria and substitution equilibria indicate that nucleotide levels are being impacted by changes in fixation probabilities due to either selection or gBGC since the species have diverged. Interestingly, the difference between equilibrium GC content as predicted by *M* and substitution rates showed a clear peak just downstream of the TSS (Figure 4E). This suggests that some evolutionary force, either selection or gBGC, has been resisting the depletion in GC-content. In addition to this peak, there was a much broader constant difference throughout the entire region which is likely due a baseline level of gBGC throughout the genome, known as background gBGC (Glémin et al., 2015).

Next, we calculated the difference between substitution equilibriums and actual GC levels (excluding CpGs) (Figure 4F). We observed a prominent peak in the difference near the TSS, with the centre of the peak again being present just downstream of the TSS. This difference means that GC content is not at the substitution equilibrium, as we previously reported (Qiu et al., 2024). This is likely due to ancient evolutionary trends that elevated GC content above its current substitution equilibrium and thus were restricted to a period before the human and chimpanzee lineages diverged. These were likely due to gBGC events near the TSS. Other groups have found that TSS-associated gBGC, likely due to localized recombination events, occur in many placental mammals (Joseph et al., 2024). However, it is possible that some selection for GC (one possible interpretation of Figure 4E) may be slightly countering this trend.

We next wanted to determine whether the increase in the fixation of GC over AT near the TSS (as seen in Figure 4E) was still ongoing and whether it was due to selection on exonic regions, where increases in GC content promotes mRNA nuclear export (Palazzo and Kang, 2021). Thus, we extracted the site frequency spectrum (SFS) of rare SNPs from gnomAD in intergenic regions and from the observed ΔGC peak around the TSS (Figure 4E, –150 to +350 around the TSS). We further parsed regions downstream of the TSS to exonic and intronic sites, as the former should be under selection while the later should not. We also examined sites of germline recombination hotspots as a positive control for gBGC. We then calculated the ratio of weak to strong (WS, SNPs towards GC) and strong to weak (SW, SNPs away from GC) alleles. One important note is that a significant number of SNPs in the gnomAD dataset are mispolarized between the ancestral and derived alleles. This polarization is negligible at low frequencies but drastically affects alleles in the higher frequency bins (Supplementary Table 3). We therefore polarized the alleles based on Ensembl’s ancestral sequence which is deduced from multi-species primate alignment (Paten et al., 2008). We observe that throughout all derived allele frequency (DAF) bins, the SW/WS ratios around the TSS is similar to that of intergenic regions. When exonic and intronic regions are examined separately they too have the same ratios. This suggests a lack of selection for GC around the TSS. Note that all ratios increase slightly (from ∼0.5 at low DAF bins to ∼0.6 at high DAF bins), consistent with background gBGC. For recombination hotspots, the ratio starts to deviate from the rest at around DAF bin 0.01 and rises drastically after DAF bin 0.2. This is a clear example of gBGC, which is not observed for sites around the TSS.

From this analysis we conclude that it is very unlikely that the increase in GC at the start of genes is due to selection, either in exons or at promoters, as we would expect this type of evolutionary force to be ongoing. Instead, the fluctuation in fixation biases towards GC is best explained by punctuated episodes of TSS-associated gBGC. Our data strongly indicates that the lineage that gave rise to the humans-chimpanzee ancestor, experienced TSS-associated gBGC episodes (Figure 4F). After humans split off from chimpanzees, our lineage experienced a limited amount of gBGC (Figure 4E). Then, in more recent evolutionary history, we see a lack of TSS-associated gBGC (Figure 4G). Since this last analysis is based in part on high-frequency human variants whose age has been estimated between 40,000 to 500,000 years ago (Andirkó et al., 2022), it is likely that there has not been any punctuated TSS-associated gBGC episodes over this timescale.

### Estimation of gBGC strength (*B*) at TSS regions

We used the polarized WS and SW polymorphism data above (Figure 4G) to estimate the population-scaled gBGC coefficient *B* for each region. We followed the diffusion-based framework of Glémin, which estimates *B* from the relative distribution of WS and SW polymorphisms across the derived allele frequency (DAF) spectrum (Glémin et al., 2015). Instead of fitting the absolute DAF, we estimated for each frequency class, the proportion of WS among SW+WS mutations. Because demography affects both WS and SW mutations similarly, much of the demographic distortion cancels in this conditional ratio, making the inference of *B* robust (Glémin et al., 2015).

To separate the contribution of gBGC from regional differences in baseline mutation rates and W vs. S base availability, we included a second parameter, *c*, which captures the expected WS:SW count ratio under *B* = 0. This parameter takes into account asymmetries introduced at the mutation stage, and differences in local base composition. Because these processes affect the creation of mutations rather than their subsequent frequency dynamics, they are expected to primarily alter the full WS:SW ratio without producing the DAF-skew generated by gBGC.

Across all regions, the intergenic estimate of *B =* 0.47 (95% CI: 0.46–0.47) reproduced previous estimates of the background gBGC (between 0.3-0.5 (Glémin et al., 2015; Galtier, 2021)), confirming the method’s calibration. Recombination hotspots gave *B* = 2.45 (95% CI: 2.40–2.50), within the expected range for human hotspots (Capra et al., 2013; Glémin et al., 2015) and consistent with the steep DAF-dependent rise visible at these regions (Figure 4G). In contrast, the TSS gave *B* = 0.33 (95% CI: 0.25–0.40), and the exon and intron sub-windows gave *B* = 0.36 and 0.22, respectively. Thus, even when we explicitly model the DAF spectrum to extract gBGC strength, the TSS region shows no excess gBGC relative to genome-wide background. Note that the value of *c* is low for TSS, consistent with high GC content at the TSS, providing less raw opportunities for WS mutations.

From this analysis we conclude that the TSS is experiencing low levels of background gBGC estimated between 0.25 and 0.40, and there is no evidence of recent (in the last ∼500,000 years, see reasoning above) TSS-associated hotspot gBGC episodes in modern humans.

### *In silico* modeling of nucleotide content based on localized mutation rates and gBGC

Previous analyses did not explicitly account for CpG dynamics, despite their potential impact on equilibrium nucleotide content estimates (Figure 4E-F). To incorporate these effects and estimate additional parameters such as TSS-associated ancestral gBGC, we developed a forward-in-time simulation algorithm. We modeled the evolution of synthetic TSS regions while incorporating different localized inferred mutation rates (including CpG decay) and gBGC. The model randomly assigns 5,000 base pairs surrounding the TSS based on the average GC-content of the human genome (42%). For each gene, it assigns several features: a nucleotide at the TSS to the midpoint (labeled 0) using the known distribution of human TSS nucleotides (Carninci et al., 2005); a 5’SS motif to an exon-intron boundary using known positional (Movassat et al., 2019) and motif (Sibley et al., 2016) distributions; and a translational start site codon, based on the distribution of 5’UTR sizes from human protein-coding genes (Leppek et al., 2018). The model then calculates a matrix of mutation rates per position for each gene by starting off with the known genome wide rate for all twelve mutations and CpGs. By fitting Gaussian curves, we then calculated the decrease in CpG mutations around the TSS using our new estimates (Figure 1D-E) and incorporated this into our matrix. Finally, we increased the base mutation rate around the TSS by fitting two Gaussian curves to match the two main peaks seen in our new estimate of the overall mutation rate at the TSS (see Figure 1B, red line). This matrix modifies the base mutation rate, which in humans is somewhere between 1.2×10^-8^ (Harris and Pritchard, 2017) to 1.88×10^-8^ (Figure 1B, black line) mutations per nucleotide. This has been set to 10^-3^ mutations per nucleotide in the simulation, indicating that each round represents 5×10^4^ to 8×10^4^ generations (assuming that the mutation rate per generation remains constant).

Several factors will then either increase or decrease the probability that a mutation fixes (i.e. fixation rate) and we incorporated this into our model by adjusting the mutation matrix, transforming it into a fixation matrix. Using estimates from ENCODE we decreased the fixation rate in the 5’UTRs by 20% and the coding region by 70% (see Figure 11B in (ENCODE Project Consortium et al., 2007)). Using our estimates of TSS-associated background gBGC (denoted as *B_b_*) (Figure 4H), we set it at 0.5 and used it to adjust all WS and SW mutation rates. We also modeled ancient increases in GC-content at the TSS due to these regions being used as recombination hotspots, as evidenced in our observation of increase gBGC events at the TSS before the human-chimp lineages diverged (Figure 4F). This results in an additional long-term ancestral *B* (denoted as *B_add_*) at the TSS due to enhanced recombination. Note that when recombination is happening at the TSS, the overall *B* in this region (denotated as *B_TSS_*) is the sum of *B_b_* and *B_add_*. We modeled this effect as a Gaussian curve with a maxima *B_TSS_* at 2.25, which is lower than PRDM9-associated hotspots (Glémin et al., 2015), but gave good preliminary results. The position of the Gaussian curve maxima was set to be located at 80 nucleotides downstream of the TSS, where GC-content is highest, and its width (or σ) was set at 300 nucleotides. This width will reflect both the average gene conversion tract length (estimated to be 75-123 nucleotides for non-crossover gene conversions, which outnumber crossover gene conversions by an order of magnitude (Williams et al., 2015; Palsson et al., 2025)) and the variability in the position of the gene conversion tract. Finally, we incorporated a time interval for how long-ago excess recombination at the TSS was turned off. At this point, *B* becomes uniform across the entire 5kb region (with *B_add_* = 0). Looking at a collection of primates, Joseph and colleagues found strong evidence for TSS-associated gBGC only in *Lemur catta* (Joseph et al., 2024), which diverged from humans approximately 63 MYA (Yoder et al., 2003). In light of this we let the simulation run for 500 rounds where there is enhanced recombination at the TSS (where *B_add_* = 1.75, and thus *B_TSS_* =2.25) and then added 100 rounds of our simulation where there was no extra recombination at the TSS and thus *B_add_* = 0. This last time interval represents about 5×10^6^ to 8×10^6^ generations. If the average generation time for primates is considered to be 6-10 years, this is equivalent to about 30 to 80 million years.

Using these parameters, we simulated the evolution of 2,000 genes and plotted their average nucleotide content. The simulation matched the observed trends well in the intergenic regions, but diverged significantly just upstream of the TSS and continuing downstream from there (Figure 5A).

**Figure 5.**
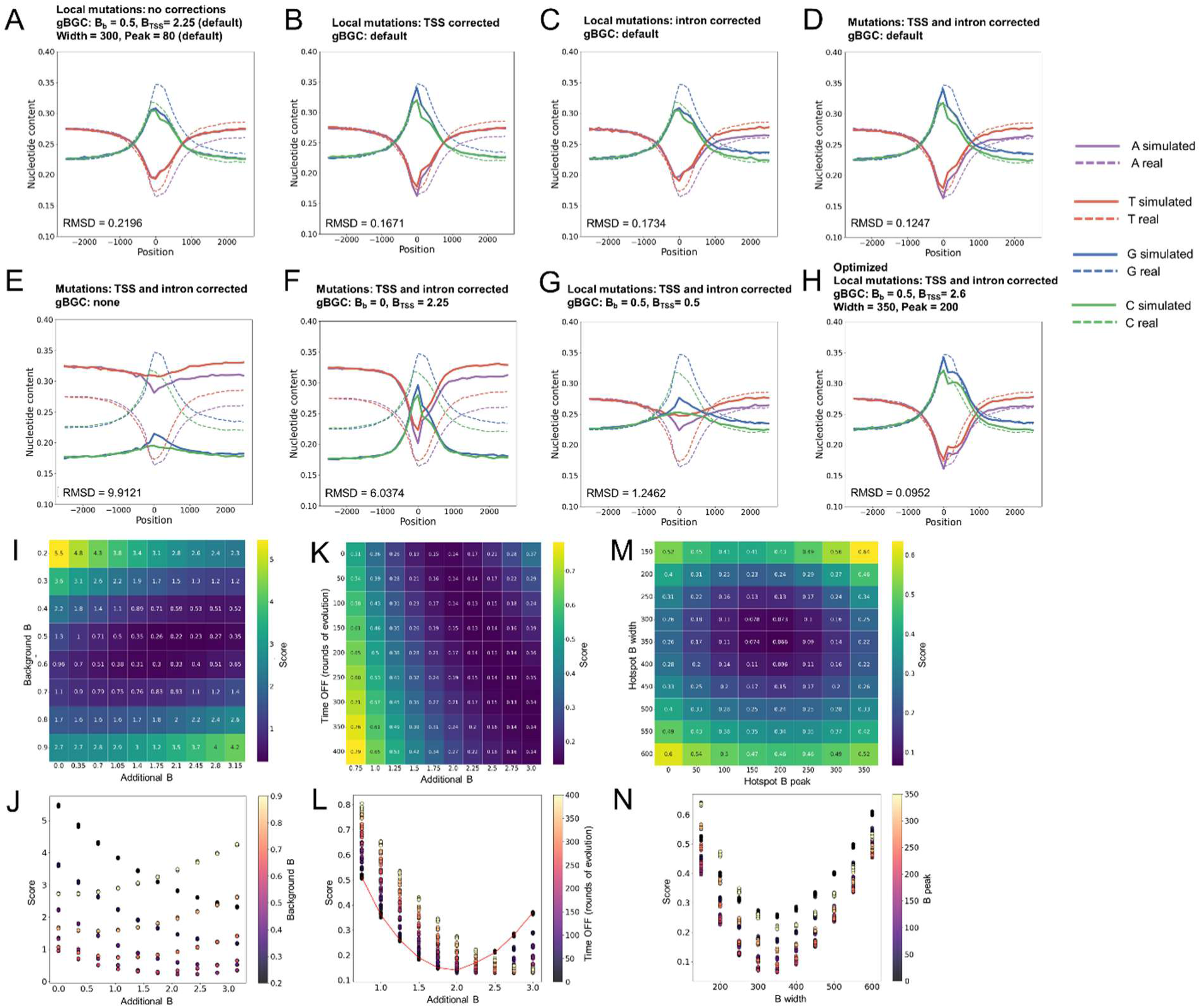
*In silico* simulations of the nucleotide content of human protein-coding genes. A-H) Results of *in silico* forward-in-time simulation algorithm to model the evolution of synthetic TSS regions. Solid lines are the average of 2,000 simulated genes. Doted lines are the average nucleotide content of all human protein-coding genes. Various parameters (local mutation rates around the TSS and/or introns, *B_b_* and *B_add_*/*B_TSS_*) were added or removed. Optimized parameters are used in (H). Root mean squared deviations (RMSDs) are indicated. I-J) Simulations were conducted with various *B_b_* and *B_add_*. For each combination, the resulting nucleotide patterns were compared to actual nucleotide values to minimize the RMSDs (scores). K-L) Simulations were conducted with various *B_add_* and rounds with *B_add_* = 0 (Time OFF). For each combination, the resulting nucleotide patterns were compared to actual nucleotide values to minimize the RMSDs (scores). Simulations in (L) with Time OFF rounds = 0, are connected by a red line plot. M-N) Simulations were conducted with various widths (σ represents one standard deviation of the normalized curve of the TSS-associated gBGC effect) and positions downstream from the TSS that experience additional recombination (“peak” represents the center of the normalized curve of the TSS-associated gBGC effect) where *B_TSS_* is applied. For each combination, the resulting nucleotide patterns were compared to actual nucleotide values to minimize the RMSDs (scores).

Next we incorporated individual inferred mutation rate corrections for nucleotides near the TSS (see Figures 1F-Q and Supplemental Figure 7, black curves). Of all individual inferred mutation rates assessed, G to A (Supplemental Figure 7D) diverged the greatest from the overall mutation rate (Supplemental Figure 7, bright red curves). We reran the simulation incorporating changes from four additional mutations that diverged the most from the overall rate. After assessing the effect of these four mutations alone or in combination, we determined that simulations that include the G to A residual rate alone (Figure 5B) had the lowest Akaike Information Criterion (AIC) among all models, indicating the best trade-off between model complexity and fit measured by likelihood (see methods).

We then incorporated inferred mutation rate corrections for nucleotides in the intron (see Supplemental Figure 3, “around exon/intron boundary” column 4, and Supplemental Figure 8). This time we reran the simulation with seven mutations (alone or in combination) and determined that including three (C to A, T to A and G to C) gave the lowest AIC. Incorporating these three corrections, without (Figure 5C) and with (Figure 5D) the TSS correction not only decreased the root mean square deviation (RMSD) but generated much of the strand asymmetry seen in the real data.

Next, we wanted to observe how much of an effect gBGC had on our simulations to assess its role in the evolution of TSS content. When all the effects of gBGC were omitted, the simulations were off by a wide margin (Figure 5E). Adding back just *B_b_* or just *B_add_*, improved the simulation a bit (Figure 5F-G), but were not as good as when both were incorporated.

We then proceeded to optimize *B* (the final simulation with optimizations is shown in Figure 5H). To do so, we reran the simulations with various *B_b_* and *B_add_*. We found that values with the lowest RMSD were *B_b_* = 0.5 and *B_add_* = 2.1 (making B_TSS_ = 2.6) (Figure 5I-J). Next, we wanted to examine the relationship between the strength of long term gBGC at the TSS and the period of time when there was no extra recombination at the TSS (Time OFF) (Figure 5K-L). Note that the data points for Time OFF = 0 rounds are highlighted (red line in Figure 5L). This line highlights that Time OFF = 0 produces a global minimum score distribution when *B_add_* < 2, and produces a maximum when *B_add_* > 2. Note that we observed that slight increases in *B_add_* would require very long periods of additional rounds without recombination to match current nucleotide levels. This likely reflects the fact that the decay in GC content that is experienced at the TSS when there is no extra recombination is very gradual. Finaly, we reran the simulations varying the width and central position of *B_TSS_* and found that the lowest RMSD scores had values where the width = 350, and the peak position = 200 nucleotides downstream from the TSS (Figure 5M-N). This was much further downstream than we had anticipated but matched our measurements of ancient gBGC (see Figure 4E-F). When we incorporated the optimized values (*B_b_* = 0.5, *B_TSS_* = 2.6, width (“σ”) of TSS associated gBGC = 350, center of TSS associated gBGC = 200, and rounds of no TSS associated gBGC = 100), this gave a simulation that closely matched the observed values for all human protein-coding genes (Figure 5H).

In summary, our *in silico* modeling suggests that biased mutations and gBGC played important roles in shaping nucleotide content around the TSS of protein-coding genes. The model predicts a *B_b_* value close to 0.5 near the TSS consistent with the estimate obtained from DAFs (0.25-0.4, Figure 4H). Our model also indicates that *B_TSS_* was centered substantially further downstream than we originally expected, but that its strength likely matched what we currently see at recombination hotspots.

## DISCUSSION

Here we provide evidence that the transcriptional start site of protein-coding genes is a hotspot for inherited *de novo* mutations and that these contribute to gene evolution. Our data suggest that these mutations are caused in part by paused RNA Polymerase II activity in germline cells (Figure 3A). Some of our data is in line with a recent study (Cortés Guzmán et al., 2025) reporting excess inherited mutations near human protein-coding TSSs. Interestingly, that study did not detect the same peak in parent-offspring trio sequencing analysis and instead attribute the increase to mutations that occurred in the germline post-zygotically. In out study, we used different parent-offspring trio sequencing data, which contained a larger number of mutations, and allowed us to detect a TSS-associated mutational peak. However, the amplitude of the mutational peak at TSSs in the parent-offspring trio sequencing data is substantially lower than what is predicted by the *M* metric, and lower than at RNA polymerase II pause sites (Figure 3C). As suggested by Cortés Guzmán et al., the difference in peak size may be due to post-zygotic mutations that are filtered out of *de novo* mutation analyses from trio-sequencing, (Cortés Guzmán et al., 2025). Beyond this finding, our analyses suggest that this mutational hotspot, along with other non-adaptive evolutionary processes, most notably gBGC, help to shape the nucleotide content around the TSS.

It is likely that other transcribed regions are subjected to elevated mutation rates. For example, it has been seen that regions transcribed by RNA Polymerase III, are also elevated in inherited *de novo* mutations (Seplyarskiy et al., 2023). It has also been observed that origins of replication impact the nucleotide content of nearby genes in an distance– and orientation-dependent manner (Cl et al., 2010; Chen et al., 2011; Seplyarskiy et al., 2019, 2021). These patterns suggest that differential replication errors and/or repair events accumulate during the replication of leading or lagging strands, a phenomenon known as R-asymmetry (Seplyarskiy and Sunyaev, 2021), and have impacts on nucleotide content of certain genes. It has also been suggested that transcription during spermatogenesis promotes DNA repair in human and mouse genes (Xia et al., 2020). Others have documented reduced mutation rates in gene bodies in *Arabidopsis thaliana* (Monroe et al., 2022), although these remain controversial (Wang et al., 2023; Liu and Zhang, 2022). Our analysis suggests the reverse in human genes, particularly the region surrounding the TSS.

Our analysis is contingent on the assumption that *M* ∼ *μ*, and that low frequency SNPs in these regions have not experienced much selective pressure. Indeed, selection should be primarily acting to purge deleterious mutations and thus suppressing variation in the low frequency SNP pool. Since most of the inferred mutation rates that we document are significantly higher than the background rate (Figure 1B, G-R), this change cannot be attributed to negative selection. Although these higher rates would be consistent with recent positive selection, this would be highly unlikely as it would require greatly altered SFS for these mutations, with a skew towards higher frequencies, and this is not seen in the data (Figure 2D).

Our data supports the notion that strand asymmetry in the nucleotide content (also known as skew) of genes can be explained by localized changes in mutational rates that are strand asymmetric (e.g. that G to A and C to T rates are different on a given strand). Our observations and simulations indicate that trends in *de novo* mutations near the TSS likely contribute to the nucleotide content of the first exons and first introns. These results may explain why introns tend to be U-rich and have extreme strand asymmetry (Palazzo et al., 2024). Our findings that differential mutations on coding and template strands may explain some of the strand asymmetry in genes, and mirror previous observations that were based on substitution rates (Polak and Arndt, 2008).

Our analyses also clarify how current patterns of recombination contributes to the nucleotide content at the TSS. The gBGC strength near the TSS (*B_b_* = 0.25 to 0.40, Figure 4H, and *B_b_* = 0.5, Figure 5H) is similar to other genome-wide estimates (between 0.3-0.5 (Glémin et al., 2015; Galtier, 2021)). Our estimation of *B_b_* in Figure 4H assumes constant effective population size, no purifying selection on W↔S polymorphisms, and stationary mutation signatures across DAF bins, which are standard population genetics assumptions of the model. Having said this, it is possible that the filtering out of CpGs may lead to slight changes in DAF ratios, thus explaining the slight discrepancy between our two estimates of *B_b_*.

Our analyses (Figure 4E-G) and simulations (Figure 5H, I-N) provide insights into how ancient recombination events shaped the GC-content of human genes. The strength of *B_TSS_*, which is a measure of the long-term gBGC strength, depends on how long ago our lineage stopped experiencing elevated recombination around TSSs (Figure 5K-L). It is likely that these ancient gBGC events were due to both crossover– and non-crossover-catalyzed gene conversion events, although the latter outnumber the former by an order of magnitude. Recent studies have begun to clarify the genome-wide distribution of non-crossover gene conversion events (Li et al., 2019; Palsson et al., 2025). In humans, they tend to correlate with recombination hotspots but are enriched in highly transcribed regions, especially in exons, beyond what is predicted by recombination rates (Palsson et al., 2025). Despite this, their relation to TSSs in various PRDM9 allele backgrounds (i.e., in species with high levels of TSS-associated gBGC) needs further investigation.

From our simulations, the region affected by *B_TSS_* is centered further downstream than we anticipated (∼200 nucleotides, Figure 5M-N). Indeed, when we compare substitution equilibria and actual GC-content, we can see that the residual, which we argue is due to gene conversion tracts from ancient recombination sites, also gives a peak that is downstream from the TSS (see Figure 4E-F). The center of this peak may still be dictated by RNA Polymerase II pausing (20 and 60 nucleotides downstream from the TSS), likely when it encounters the first nucleosome (Naganuma et al., 2025). One possible explanation is that transcriptional bubbles generate supercoiling that is relieved by topoisomerases acting downstream from paused RNA Polymerase II. Indeed, topoisomerases induce double-stranded breaks in these regions (Singh et al., 2020), and in particular, SPO11 initiates the double-stranded break at PRDM9 sites.

Our models predict that the width of *B_TSS_* = 350 nucleotides (the σ of a gaussian curve, Figure 5M-N). This peak width is a combination of gene conversion lengths, estimated to be 75-123 nucleotides long (Williams et al., 2015; Palsson et al., 2025), and the variability in the central position of gene conversion tracts. These likely vary between genes, and between individual events at the same gene. When compared to PRDM9 motifs, the center of non-crossover conversion tracts have a variance of 50-100 nucleotides (see Figure S6 in (Palsson et al., 2025)). If we assume a similar variability in the center of the tracks, this combined with track length may explain our overall width.

The non-adaptive sculpting of genomic features may represent an important mechanism underlying the emergence and the maintenance of genomic landmarks in organisms that evolve under weak selection regimes, such as humans. In the absence of these non-adaptive forces, selection alone may be insufficient to maintain many of these landmarks. Instead, selection may primarly act on molecular processes, such as those involved in the mRNA metabolism, to recognize and use these genomic patterns that are largely shaped by non-adaptive evolutionary forces.

## DATA AVAILABILITY

All python codes can be obtained from the online repository: https://github.com/tinaqiu221/TSS_hypermutation

Some references were made to codes deposited for a previous publication (Qiu et al., 2024): https://github.com/tinaqiu221/GC_evolution

All raw data can be obtained from the online repository: https://doi.org/10.5281/zenodo.20292802

## ACKNOWLEDGEMEMTS

We would like to thank Laurent Duret for insightful conversations and feedback on the manuscript. This work was funded by a grant to A.F.P. from the Natural Sciences and Engineering Research Council of Canada (FN 492860) and the Jean D’Alembert Fellowship program (France 2030 program ANR-11-IDEX-0003).

## METHODS

### Sequence data and annotation

Sequences of Human (*Homo sapian*, GRCh38, release 108), Chimpanzee (*Pan troglodytes*, Pan_tro_3.0, release 108), and Gorilla (*Gorilla gorilla*, gorGor4, release 108) genomes were accessed and downloaded from the Ensembl database (www.ensembl.org). Annotations for protein-coding genes were retrieved using the University of Santa Cruz (UCSC) Genome Browser annotation track database based on the corresponding genome assemblies. For mapping rare SNPs and *de novo* mutations, annotations for the TSS were obtained from testis tissue-specific Cap Analysis of Gene Expression (CAGE) sequencing from the FANTOM5 project (Noguchi et al., 2017). The best TSS from CAGE-seq was defined as the transcript with the highest tags per million score. This set of testes-specific CAGE-seq annotated genes was used as our default set of “protein coding genes” unless stated otherwise. For mapping substitutions between species, positions for the TSS were determined by the most commonly used start site of all transcripts per gene based on Ensembl annotation as described previously in (Qiu et al., 2024). The exon/intron boundaries corresponding to the TSSs were obtained from Gene Transfer Format annotation files from Ensembl (https://ftp.ensembl.org/pub/release-108/gtf/homo_sapiens/)

Human lncRNAs were obtained from FANTOM CAT (Hon et al., 2017). Testis samples were identified within the FANTOM-CAT gene-level expression atlas by cross-referencing sample library IDs (CNhs codes) against the FANTOM5 human tissue sample metadata file (HumanSamples2.0 SDRF), retaining columns whose sample description contained the keyword “testis.” For lncRNAs not expressed in testis, genes with Counts Per Million of over 0 in any of the three adult testis samples were excluded.

RNA Polymerase II pause sites were obtained from ENCODE ChIP-seq (https://www.encodeproject.org/experiments/ENCSR803FAP/; GEO:GSE105816), and filtered for pause sites in intergenic regions. We used the middle of the annotated ChIP-seq coverage to align each site.

To filter for genes not expressed in testes, we used gene annotations from Ensembl as described above instead of CAGE-seq. Tissue-specific RNA transcript expression was obtained from the Human Protein Atlas (https://www.proteinatlas.org/humanproteome/twassue/data#consensus_twassues_rna) (Thul and Lindskog, 2018), and only protein-coding genes with transcript per million (TPM) of 0 were included.

Intergenic genomic regions were obtained by randomly generating genome coordinates outside of protein-coding regions. (Github: tinaqiu221/GC_evolution/generate_random_coordinates.py).

Recombination hotspots were identified by intersecting two complementary genome-wide datasets. Meiotic double-strand break (DSB) hotspot coordinates were obtained from (Pratto et al., 2014), who mapped DSB initiation sites genome-wide in human testis tissue using DMC1 chromatin immunoprecipitation followed by single-stranded DNA sequencing (SSDS). Peak coordinates for PRDM9-A homozygous individuals (AA1 and AA2), representing the most common PRDM9 allele in European-ancestry populations, were downloaded from the Gene Expression Omnibus (GEO: GSE59836; GSE59836_Peak_data_Supplementary_File_1.txt.gz). Hotspot coordinates were originally mapped to hg19 and lifted over to GRCh38 using the UCSC liftOver utility with the hg19ToHg38 chain file (https://hgdownload.soe.ucsc.edu/goldenPath/hg19/liftOver/). Crossover rate data were retrieved from the UCSC Genome Browser recombAvg track via the UCSC REST API (https://api.genome.ucsc.edu), and regions with a recombination rate of ≥50 cM/Mb were retained as crossover hotspots, corresponding to approximately 40-fold above the genome-wide average. Intervals narrower than 500 bp were excluded as bigWig boundary artifacts. Final high-confidence germline recombination hotspots were defined as the intersection of DMC1-SSDS DSB peaks with crossover hotspot intervals using BEDTools intersect (Quinlan and Hall, 2010). (Github: tinaqiu221/TSS_hypermutation/recombination_hotspots_pipeline.sh).

Nucleotide sequences from all desired regions were retrieved using the BEDTools *getfasta* suite (https://bedtools.readthedocs.io/en/latest/content/tools/getfasta.html)

### Rare SNP mapping and derivation of M

Human single nucleotide polymorphisms (SNPs) were obtained from the gnomAD v3.0 database, which contains of 71,702 whole genome sequences across 9 population groups mapped to the GRCh38 genome build (https://gnomad.broadinstitute.org/news/2019-10-gnomad-v3-0/). SNPs with a frequency of less than 1% (for *M_1%_*) or less than 0.1% (for *M_0.1%_*) in all subpopulation groups was filtered using Hail python package (https://pypi.org/project/hail/). Rare SNPs were mapped 1kb around TSSs of protein-coding genes, from the front of exon 1, from the back of exon 1, around first exon/intron boundaries, around TSSs of lncRNAs expressed in testes, around TSSs of lncRNAs not expressed in testes, around intergenic RNA Polymerase II pause sites, around TSSs of protein coding genes not expressed in testes, and around random intergenic regions. The summary mutation table reports the counts of the 12 mutation types, with each count corresponding to the sum of occurrences within a 100 bp window upstream (−400 to –300 relative position), around TSS or focal site (−50 to +50 relative position), and downstream (+300 to +400 relative position).

The fraction of nucleotides with SNP frequency below the cutoff *x* (*F_x_*) were calculated by dividing the total count of rare SNPs per position by twice the number of nucleotides analyzed per relative position (to account for diploidy). In most cases, all mutations were considered; however to calculate *B* (i.e. Supplemental Figure 5, and Figure 5), CpG mutations were filtered out by excluding CG>TG and CG>CA mutations from the numerator and CG positions from the denominator for C to T and G to A mutation types. To account for the fact that multiple mutations at the same site are independent and can lead to underestimation of the true rare SNP fraction (*F^t^*), we transformed *F*:

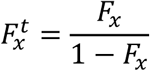

*F^t^* and genome sample number (*N*) were used to obtain the metric *D_x_* (population-scaled SNP density per site, for bin *x*):

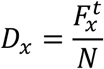

*D_x_* can be interpreted as the probability that a position contains a recently created *de novo* mutation (i.e., a rare SNP less than frequency *x*) that is observable in one member of the population. At equilibrium, and assuming limited temporal changes in parameters (*N_e_*, *s*, and *b*), the number of SNPs gained in a region of length *L* (that are in bin 0 to *x* frequency) in the surveyed population (*N*) must equal to the number of SNPs in this same region that are lost from the bin in this population. Assuming that the number of SNPs gained due to *de novo* mutations in a generation is vastly greater than the number of higher frequency SNPs that drop below *x* frequency, this gives:

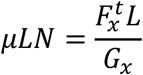

which simplifies to:

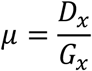

In this equation, *G_x_* is the mean number of generations a SNP is present in the ascribed bin (from 0 to *x* frequency) before being lost. It also acts as a rescaling factor between *μ* and *D_x_*. Note that *G* will depend on local *N_e_*, *s* and *b*. Using *μ* and *D* values for intergenic regions, we can estimate *G* intergenic (*G^I^*) to be 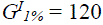 generations and 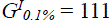 generations. Also note that our calculated intergenic mutation rate (1.88×10^-8^ mutations/base) is equivalent to 124 mutations per generation per diploid genome.

For regions such as the TSS, we can use local values of *D* (in this case *D_x_^T^*) to obtain a metric *M_x_^T^*:

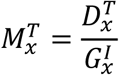

Assuming that *G* is the same in intergenic and TSS regions (i.e., that *G^I^* ≈ *G^T^*), which is true when *s* and *b* are small and local *N_e_* in these two regions are similar, then *M^T^* ≈ μ. Unless stated otherwise, *M* refers to *M_1%_*.

In calculations of the fractional change between *M_1%_* and *M_0.1%_*, CpG mutations were omitted as these significantly contribute additional SW mutations that exaggerate the fractional change due to gBGC effects (an increase in SW mutations to *M_1%_* and depletion to *M_0.1%_* as can be seen in Supplemental Figure 8).

All codes were compiled and annotated in Jupyter Notebook (Github: tinaqiu221/TSS_hypermutation/RareSNP.ipynb).

### 5-mer sequence context rate calculation

Overall baseline mutation rates for each nucleotide were first calculated using rare SNPs mapped to random intergenic regions. The intergenic sequences were then scanned to count all possible 5-mer contexts (the centre base flanked by two upstream and two downstream nucleotides), providing background context frequencies. For each 5-mer, a context-specific change value was computed as the context mutation rate divided by baseline rate for that nucleotide, representing the relative mutability of each context compared to the genome-wide average. The same procedure was applied to rare SNPs mapped to 1kb around regions of interest (TSSs of protein-coding genes, TSSs of lncRNAs expressed in testes, TSSs of lncRNAs not expressed in testes, intergenic RNA Polymerase II pause sites, TSSs of protein coding genes not expressed in testes). Mutations were assigned to positions relative to the feature anchor (e.g. TSS), and a 1024 × 1001 count matrix was constructed (5-mer contexts × positions). Each count was weighted by its corresponding change value. At each position, the weighted sum was divided by the total raw count to produce a context-corrected rate. This rate was multiplied by the intergenic base mutation rate to give the absolute expected mutation rate per position, reflecting the contribution of local sequence composition to mutational patterns across the region. 5-mer context rates were also calculated excluding positions and mutations in CpG contexts.

All codes were compiled and annotated in Jupyter otebook (Github: tinaqiu221/TSS_hypermutation/5mer_context_rates.ipynb

### Single generation de novo mutation mapping

Human *de novo* mutations from parent-offspring trios were obtained from Decode (Palsson et al., 2025) and mapped to 1kb regions around either TSSs of protein-coding genes, TSSs of lncRNAs, intergenic RNA Polymerase II pause sites, TSSs of protein genes not expressed in testes, and random intergenic regions. To validate results, *de novo* mutations from a pooled resource collected by Seplyarskiy et al. (Seplyarskiy et al., 2023) were mapped to 1kb regions around TSSs of protein-coding genes and similarly analyzed. Summary mutations per region and mutation rates were calculated similarly to the above rare SNPs mapping. All codes were compiled and annotated in Jupyter Notebook (Github: tinaqiu221/TSS_hypermutation/DNM.ipynb).

### Somatic mutation mapping

Somatic mutations from normal human tissues were obtained from SomaMutDB 2.0, which aggregates whole-genome sequencing data from 47 published studies encompassing 30 tissue types (Shea et al., 2026). Nine datasets involving tissues with known DNA repair deficiencies, disease conditions, or exogenous mutagen exposure (including Cockayne syndrome and xeroderma pigmentosum neurons, ulcerative colitis and IBD-affected colon, cirrhotic liver, BRCA1/2 carriers, and treatment-exposed samples) were excluded, leaving mutations from healthy tissues across 38 studies. Somatic mutations were mapped 1kb around the TSS of protein coding genes. Outlier loci contributing disproportionate mutations at individual positions were identified and excluded. All codes were compiled and annotated in Jupyter Notebook (Github: tinaqiu221/TSS_hypermutation/Somatic_mutations.ipynb)

### Nucleotide substitution mapping

For protein-coding genes, substitutions between species were obtained as described previously in (Qiu et al., 2024). (Github: tinaqiu221/GC_evoluation/get_homologous_sequences.py.ipynb, tinaqiu221/GC_evoluation/needleman_alignment.py).

### Equilibrium nucleotide content calculation

Predicted equilibrium nucleotide frequencies were computed from rare SNP *M_1%_*, *de novo* mutation rates, or substitution rates, divided into 40 bins (each containing 25 positions). All rates are calculated by dividing mutational count by mutational opportunities. For each genomic bin, a 4×4 rate matrix Q was constructed, with diagonal elements set so rows summed to zero. The equilibrium distribution π was obtained by solving πQ = 0 using least-squares optimization, subject to Σπ = 1. Predicted equilibrium frequencies were compared against actual nucleotide compositions across 40 bins spanning 5kb or 1kb positions around the TSS. Mutations in a CpG context were parsed out. GC difference was calculated either by subtracting the average of human and chimp predicted equilibrium GC based substitutions rates by the predicted equilibrium GC content based on *M_1%_*, or by subtracting the actual GC content by the average of human and chimp predicted equilibrium GC based on substitutions rates. GC differences use a sliding window of 125 nucleotides.

All codes were compiled and annotated in Jupyter Notebook (Github: tinaqiu221/TSS_hypermutation/Equilibrium.ipynb).

### Site frequency spectrum analyses

Single-nucleotide variants and global allele frequencies were extracted from the gnomAD v3.1.1 sites Hail Table for three genomic region sets: a −150 to +350 bp around the TSS of protein coding genes, deCODE-defined recombination hotspots, and a control set of random intergenic regions. Variants downstream of the TSS window were further classified as exonic or intronic using GENCODE v40 exon coordinates.

Alleles were polarized using the Ensembl EPO 6-primate inferred ancestral sequence (Paten et al., 2008), restricting to high-confidence ancestral calls. For each variant, the derived allele frequency (DAF) was computed as the alt-AF when the reference matched the ancestor, or 1 − alt-AF when the alternate matched the ancestor. Variants with no ancestral call or with an ancestor matching neither allele (recurrent / triallelic) were excluded. Polarized SNPs were classified as weak-to-strong (WS: A to C, A to G, T to C, T to G) or strong-to-weak (SW: G to A, G to T, C to A, C to T) substitutions and binned by DAF.

To account for differences in mutational opportunity across regions of differing GC content, the WS/SW ratio in each DAF bin was multiplied by the regional ratio of strong (G+C) to weak (A+T) base counts (n_S/n_W). The pipeline outputs, for each region, the GC-normalized WS/SW ratio across log-spaced DAF bins from 10⁻⁴ to 1.

All codes were compiled and annotated in Jupyter Notebook (Github: tinaqiu221/TSS_hypermutation/SFS.ipynb).

### Estimation of B

For each region, we estimated the population-scaled gBGC coefficient *B* = 4*bN*_e_, where *Nₑ* is the effective population size and *b* is the per-generation conversion bias toward strong (G/C) alleles. We followed the Wright-Fisher diffusion-based framework of Glémin et al. (2015), in which the derived allele frequency (DAF) distribution under gBGC strength *B* follows

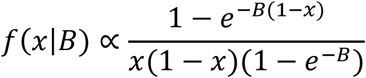

for WS variants, with *B* replaced by *-B* for SW variants (Muyle et al., 2011).

To minimize the impact of human demography which strongly distorts the SFS due to recent population expansion (Bergström et al., 2021), we did not fit *B* from the absolute SFS. Instead, we estimated the WS proportion among WS+SW for each DAF class. This proportion depends only on *B* because demography affects WS and SW spectra symmetrically and cancels out in the ratio (Glémin et al., 2015):

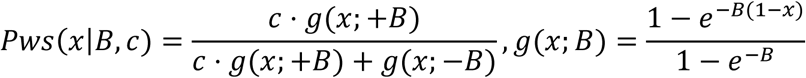

The auxiliary parameter *c* is a region-specific baseline that captures the expected raw WS:SW count ratio under *B* = 0, encompassing per-base mutation rates, W versus S base availability, and any region-specific mutation-rate effects.

For each region, polarized W↔S polymorphisms were binned into 30 log-spaced DAF intervals over [10⁻⁴, 0.95], excluding the rare extreme to avoid singleton/sequencing noise and the fixed-allele pile-up that violates infinite-sites assumptions. Within each bin *i*, observed counts were modeled as Binomial. The joint maximum-likelihood estimate of (*B*, *c*) was found by Nelder-Mead optimization, with 95% confidence intervals on *B* obtained from profile likelihood (cutoff Δ log L = 1.92, corresponding to χ²₁ at α = 0.05).

All codes were compiled and annotated in Jupyter Notebook (Github: tinaqiu221/TSS_hypermutation/B_estimation.ipynb).

### In silico modeling of nucleotide content

We developed a forward-in-time nucleotide evolution simulator to model mutation patterns around human TSS. We simulated a sequence of 5kb centered around TSS that mimics human genes that have exon and introns. We thus estimated the size of the first exon of our simulated sequences from the distribution of observed exon lengths. For each set of simulation parameters we simulated 2000 genes. We performed 10 replicates per set of parameters. We integrated 43 parameters from empirical mutation data deduced from gnomAD, including baseline mutation rates, position-specific CpG effects, and regional constraints around exons, introns, and UTRs. We also incorporated GC-biased gene conversion, with both a uniform background intensity and a localized hotspot effect, as well as a phase in which hotspot activity ceases. We evaluated models with and without a position-dependent mutation spectrum around the TSS. Parameter values were optimized to reproduce the observed per-position nucleotide frequencies, and model selection was guided by likelihood-based criteria. Using this framework, we show that both an ancestral hotspot *B* and a mutation spectrum varying along the TSS are necessary to reproduce the empirical GC content patterns. Details of the simulator, parameterization, and fitting procedure are provided in Supplementary Text 1.

(Github: tinaqiu221/TSS_hypermutation/Nucleotide_evolution_simulator/).

### COSMIC SBS mutational signatures

Mutational signature activities were analyzed across the 1kb region surrounding each genomic region in 50bp windows, yielding 20 windows per region. For each window, single base substitution (SBS) mutational signatures were assigned by fitting the observed SBS96 mutation spectrum against COSMIC v3.5 reference signatures (GRCh38 genome build) using non-negative least squares (NNLS). Signatures contributing less than 5% of total mutations in a given window were excluded. Two versions of the analysis were performed: one fitting raw mutation counts directly, and one normalizing for trinucleotide opportunity by dividing both the observed spectrum and COSMIC reference profiles by the trinucleotide composition of each 50bp window, derived from the reference genome sequence at the corresponding coordinates. Signature activities are reported as the percentage of total mutations attributed to each signature per window. All analyses were performed in Python using SigProfilerMatrixGenerator for trinucleotide context extraction, scipy for NNLS fitting, and are fully annotated in the accompanying Jupyter notebook. (Github: tinaqiu221/TSS_hypermutation/COSMIC_SBS.ipynb).

### Statistical analyses

To statistically test whether *M* or *de novo* mutation rate at one locus of a sequence was significantly different from *M* at another locus of the same sequence (e.g. between position –400 to –300 and position –50 to +50 relative to the TSS), the Wilcoxon signed-rank test was performed between the distributions of the 2 loci (Github: tinaqiu221/TSS_hypermutation/Stats_test.ipynb – Wilcoxon).

To statistically test whether the number of net change in rare SNPs or DNMs at relative nucleotide position 0 was significantly different between two datasets (e.g. between the TSS and random intergenic position), a permutation test was performed (Wilcox, 2022). In this test, net change in rare SNPs or DNMs from datasets A and B were randomly shuffled into two groups 1000 times, and the differences between the two random groups from the 1000 randomizations were used to generate a normalized curve. The actual observed delta between datasets A and group B was plotted onto the normalized curve, and the p-value was the area under the curve separated by the line x=observed delta (Github: tinaqiu221/TSS_hypermutation/Stats_test.ipynb – Permutation test). To represent this data in the figures, the normalized curve was plotted as a whisker-bar graph and the observed delta was plotted as a black dot on the same graph.

## Figures and Figure Legends

**Supplemental Figure 1.**
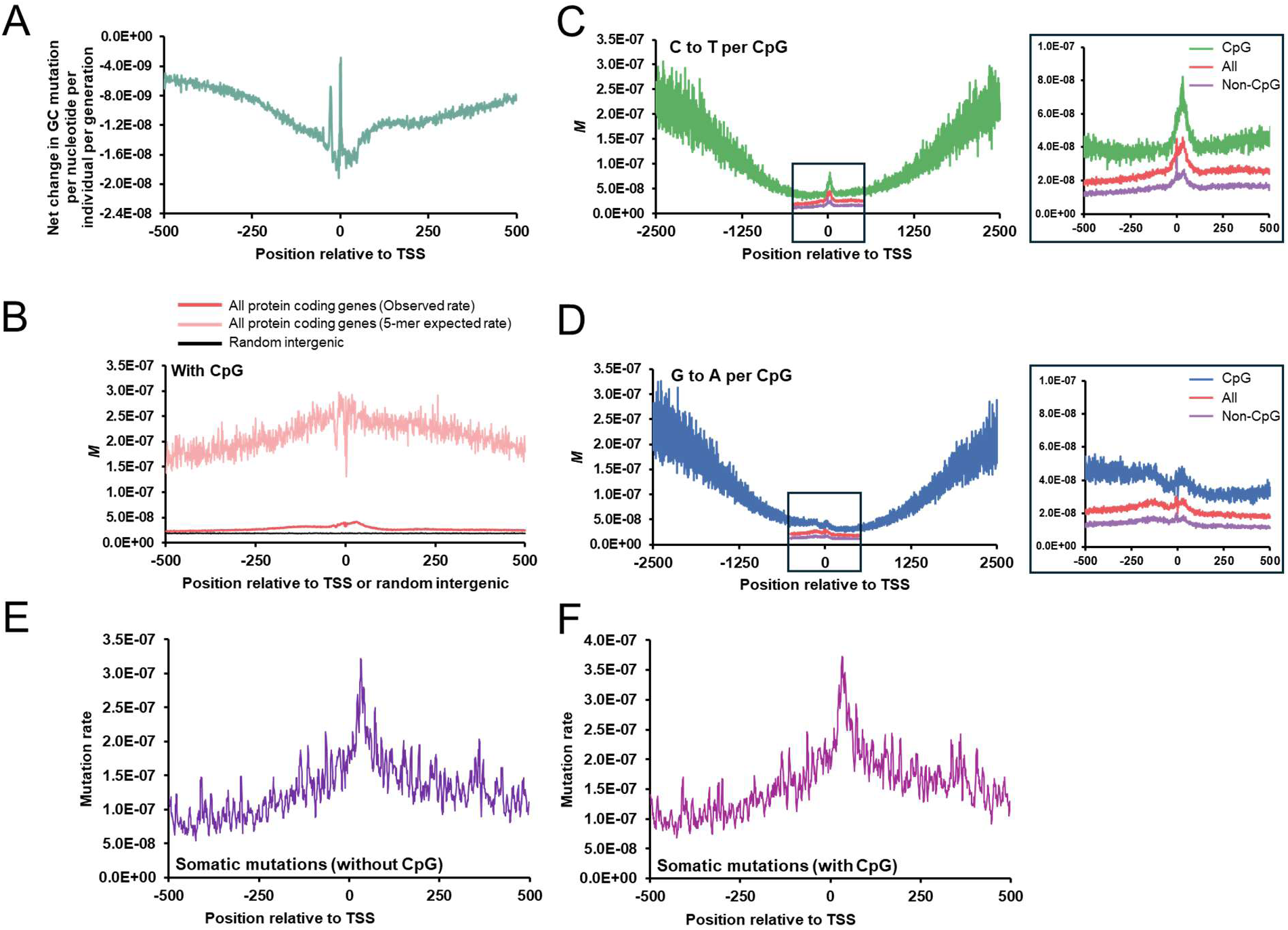
Mutation patterns around TSSs of human protein-coding genes. A) Single nucleotide resolution of *M_1%_* for ΔGC (change in GC/all countable nucleotides) around the TSS of all human protein-coding genes. Change in GC is all gain of GC mutations minus all loss of GC mutations. B) Single nucleotide resolution of *M_1%_* for all mutations including CpGs, over all TSSs from human protein-coding genes (red) or an equivalent number of intergenic regions (black) using low frequency SNP analysis from gnomAD. Single nucleotide predicted rate based on 5-mer sequence context over all TSSs from human protein-coding genes (pink). C-D) Single nucleotide resolution of *M_1%_*, for CpG to TpG (C) and CpG to CpA (D) around the TSS of all human protein-coding genes. E-F) Somatic mutations around the TSS of protein coding genes based on the SomaMutDB excluding CpG mutations (E) and including CpG mutations (F).

**Supplemental Figure 2.**
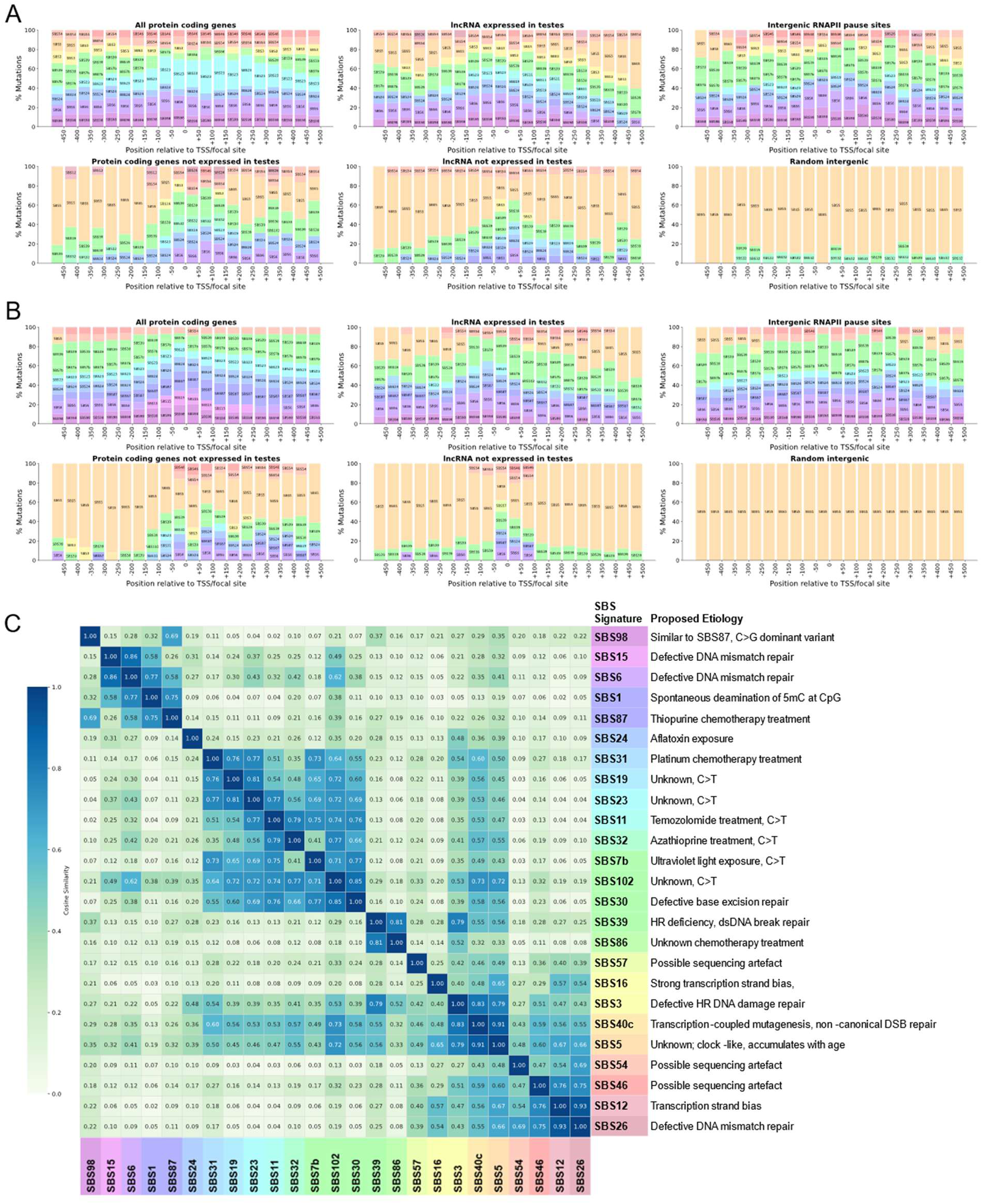
Single base substitution mutational signature activities across genomic regions. A-B) SBS mutational signature activities across 50bp windows spanning the 1kb of various genomic regions, expressed as percentage of total mutations per window, normalized for trinucleotide opportunity (A) or not normalized (B) C) Pairwise cosine similarity between COSMIC v3.5 SBS reference profiles for all signatures identified as active in (A-B)

**Supplemental Figure 3.**
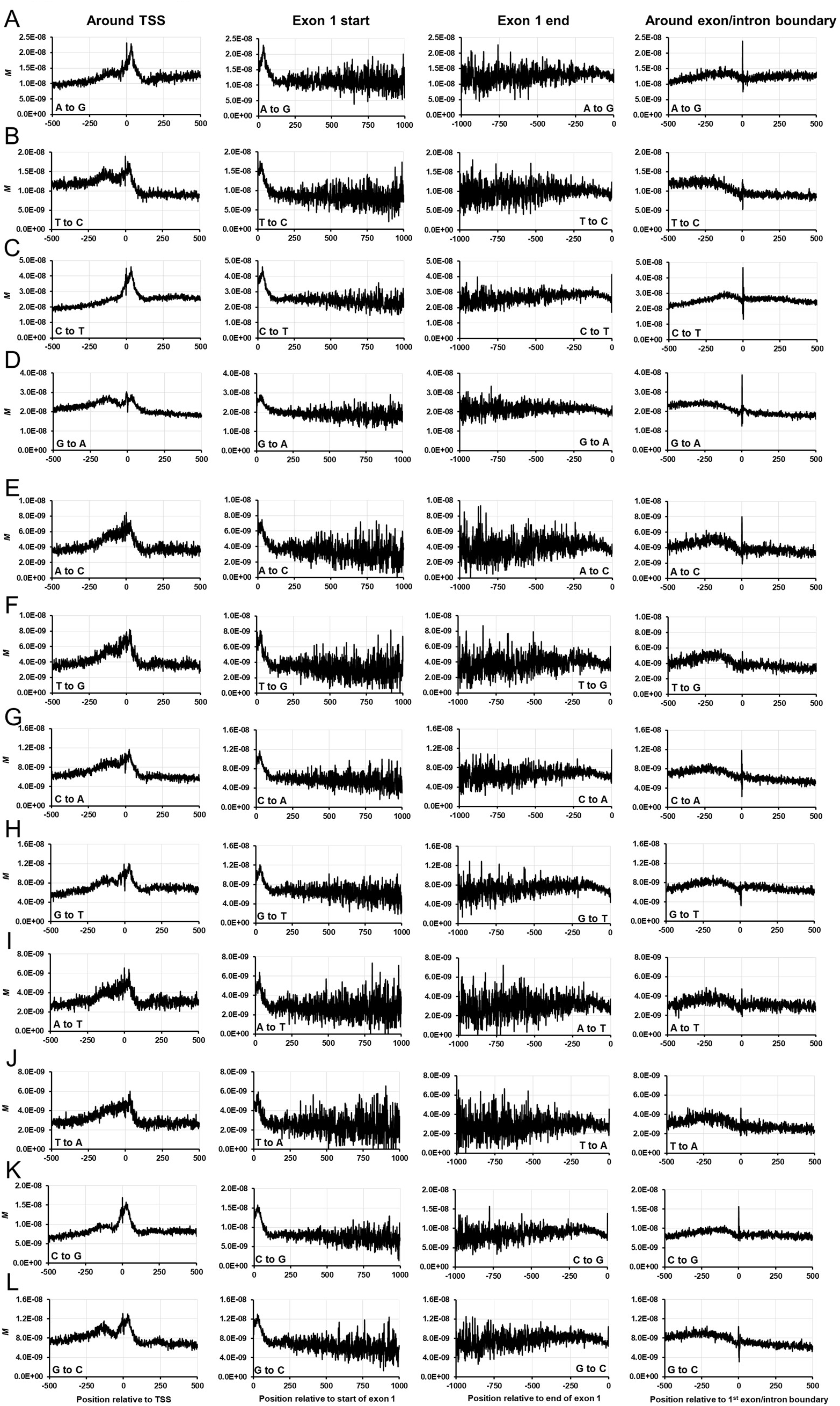
Single nucleotide resolution *M* for the first exon of human protein-coding genes. A-L) Single nucleotide resolution of *M_1%_*, for each indicated mutation type around the TSS (column 1) and first exon/intron boundary (column 4) of all human protein-coding genes. Column 2 is all first exons aligned so that the first nucleotides are positioned at 0. Column 3 is all first exons aligned so that the last nucleotides (at the exon/intron boundary) are positioned at 0.

**Supplemental Figure 4.**
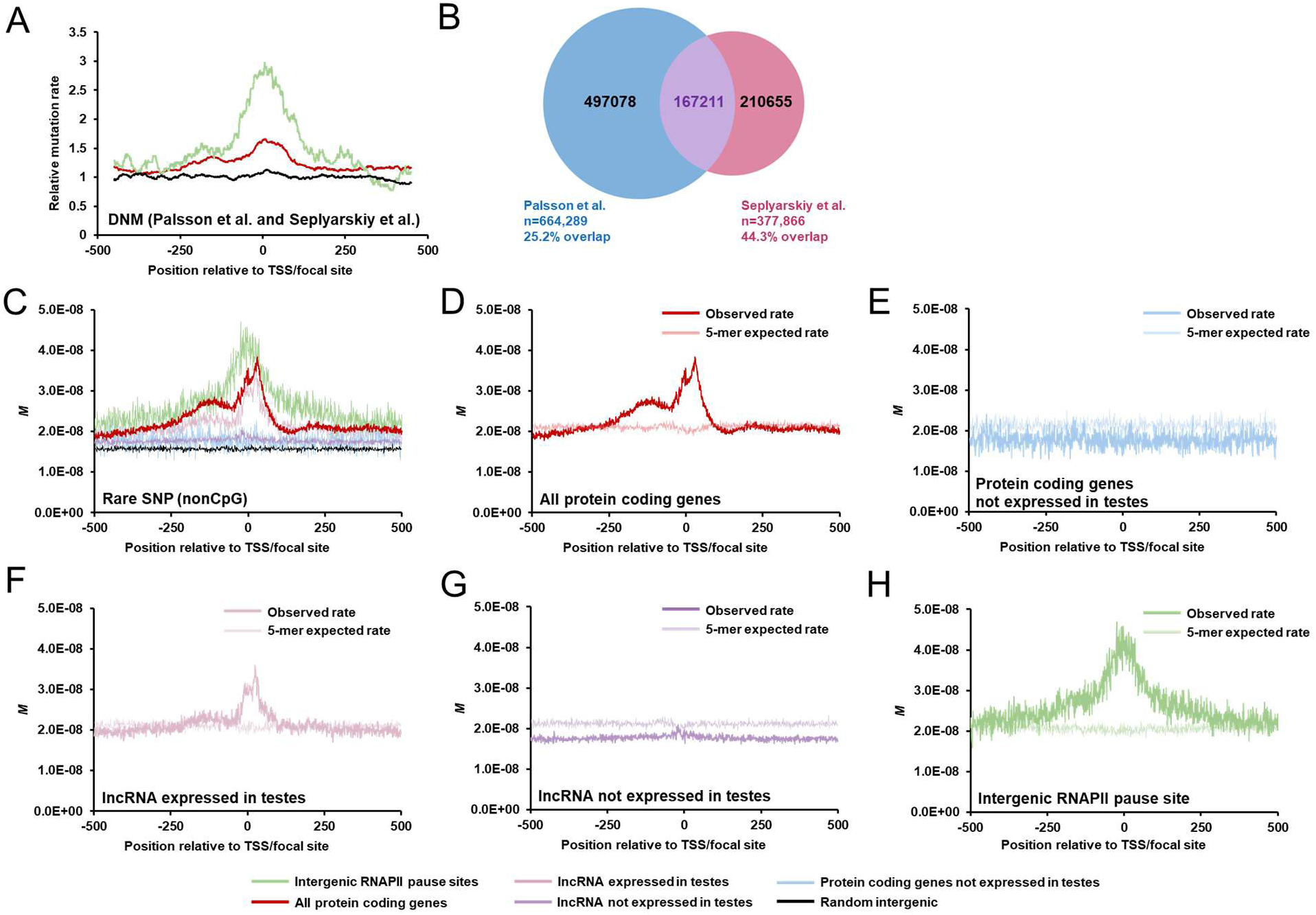
Mutation rate analyses across genomic regions. A) Mutation rates, averaged over all indicated regions from the human genome, based on *de novo* mutations as inferred by parent-offspring trio sequence analysis. Rates are plotted as a sliding window of 100 nucleotides. B) Number of DNMs from Palsson et al. and Seplyarskiy et al. datasets, and overlapping counts. C) Single nucleotide resolution of *M_1%_* for all mutations excluding CpGs, over various indicated regions from the human genome. D-H) Single nucleotide resolution of *M_1%_* for all mutations excluding CpGs, over all TSSs from human protein-coding genes (D), protein-coding genes not expressed in testes (E), lncRNA expressed in testes (F), lncRNA not expressed in testes (G), and intergenic RNAPII pause sites (H), using low frequency SNP analysis from gnomAD. Single nucleotide predicted rate based on 5-mer sequence context over regions indicated are in lighter shaded lines.

**Supplemental Figure 5.**
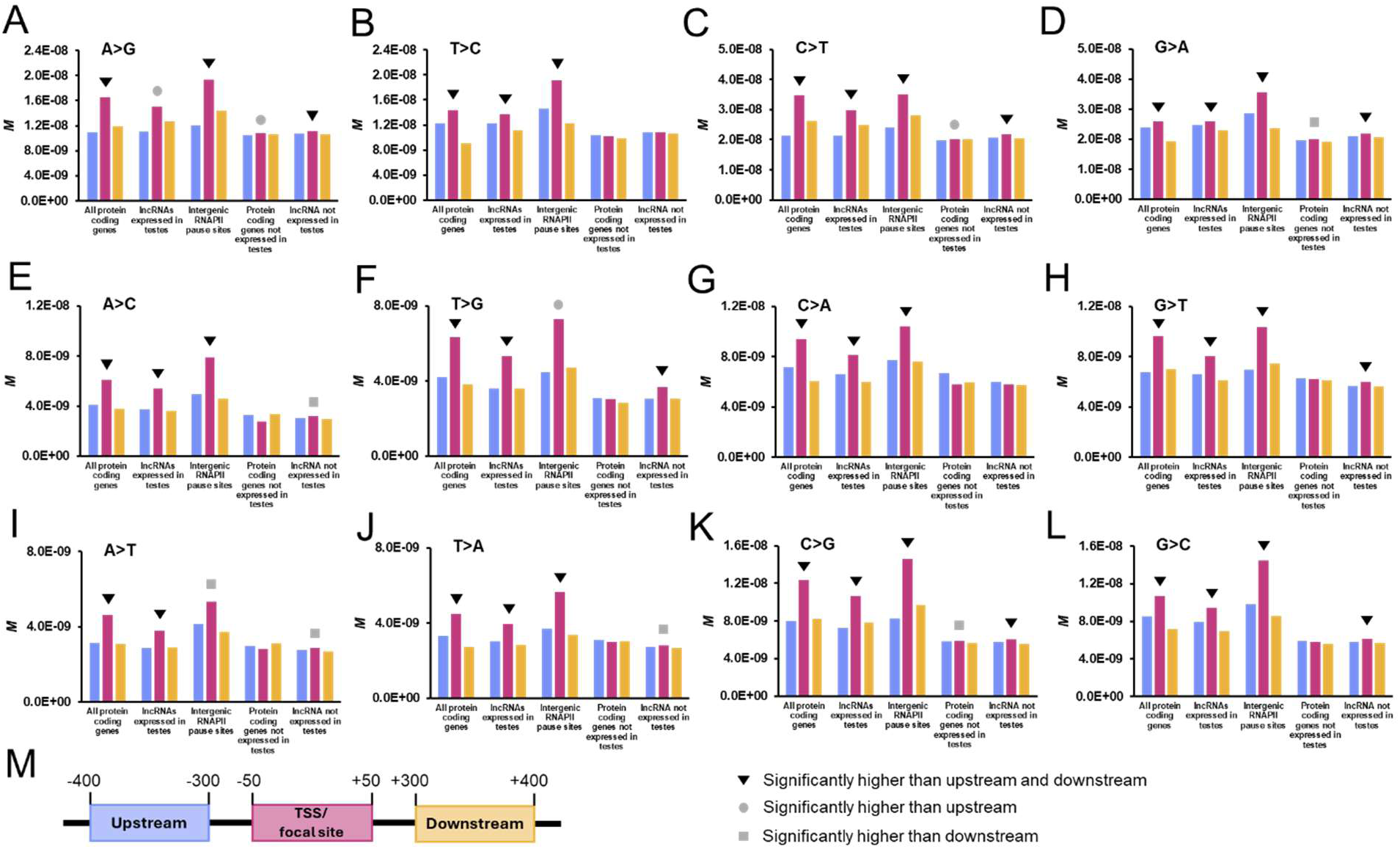
*M* of individual mutation types in various genomic regions. A-M) *M_1%_* over 100 bp windows as indicated in (M) for various indicated regions from the human genome. Statistical significance (p < 0.05) results from Wilcoxon signed-rank tests are indicated in symbols.

**Supplemental Figure 6.**
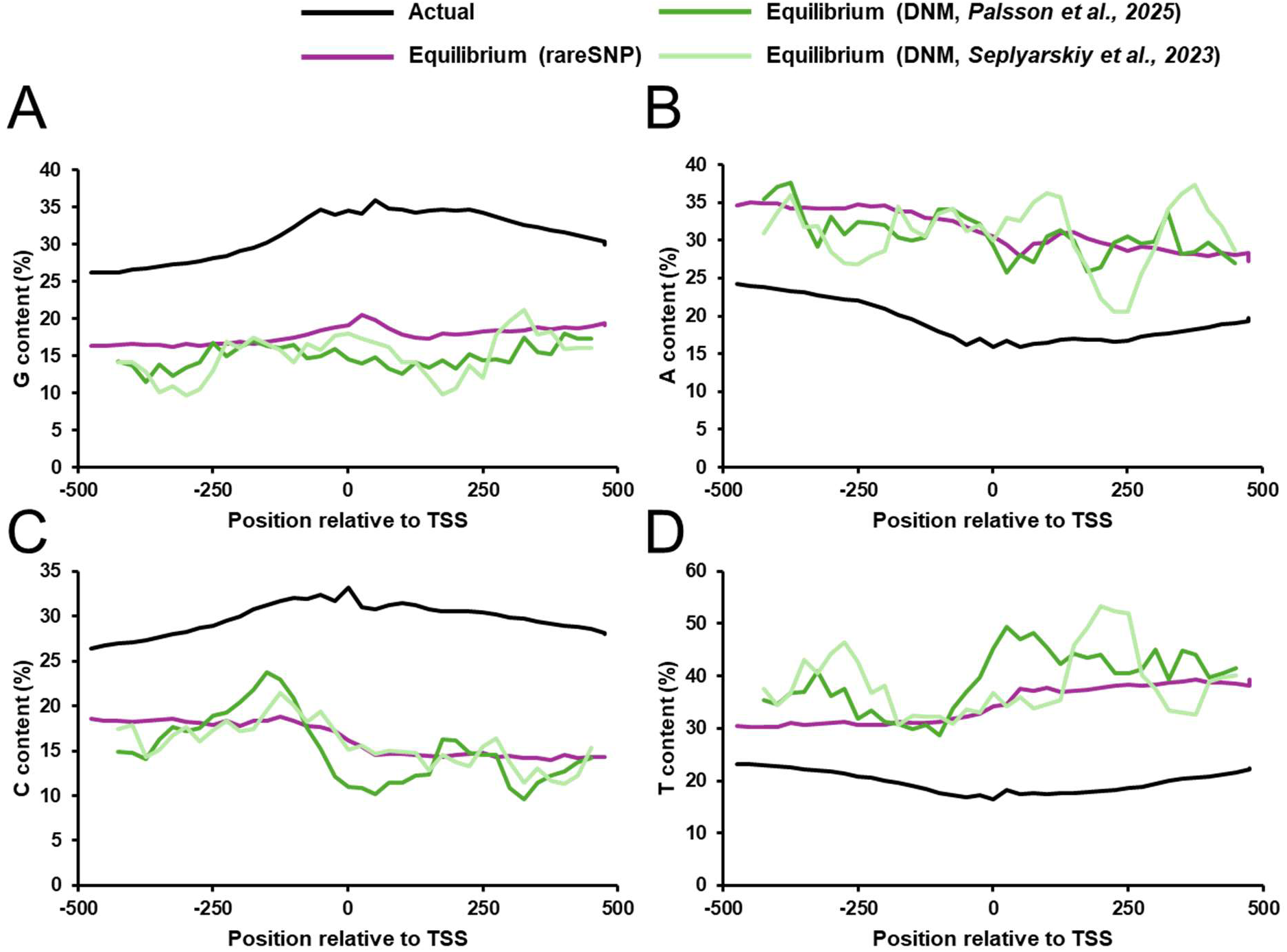
Calculated equilibrium levels of nucleotides inferred from *M* and *de novo* mutation rates. A-D) The observed (black lines) nucleotide content and predicted equilibrium nucleotide levels based on either *M_1%_* (purple lines) or *de novo* mutations as inferred from parent-offspring trio sequencing from Decode (Palsson et al., 2025) (dark green) or a list compiled by a second study (Seplyarskiy et al., 2023) (light green). Observed nucleotide content and equilibrium predicted based on *M_1%_* are in bins of 25 nucleotides, and equilibrium predicted based on both *de novo* mutations are in sliding window of 125 nucleotides.

**Supplemental Figure 7.**
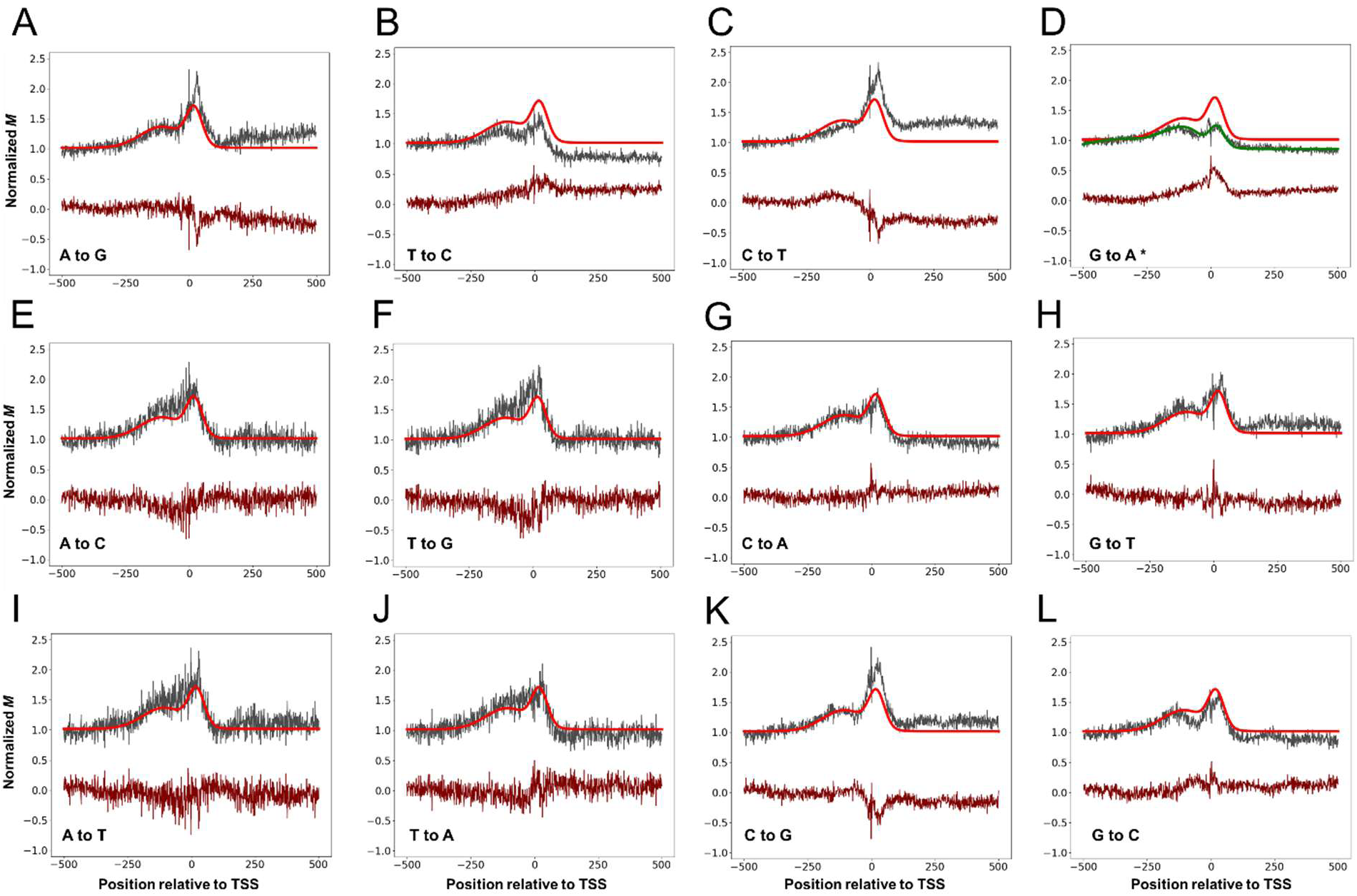
Comparison of overall *M* at TSS with individual rates. A-L) Normalized inferred mutation rates *M* (grey lines) are compared to line fitted overall inferred mutation rate *M* used in the *in silico* time-forward simulation of nucleotide content evolution (smooth bright red line). The resulting residual (burgundy line) is also plotted. After AIC analysis, it was determined that incorporating the G to A residual (D) significantly improved the simulation performance and the resulting mutation rate (fitted overall mutation rate + fitted residual) is also plotted (smooth green line) in (D).

**Supplemental Figure 8.**
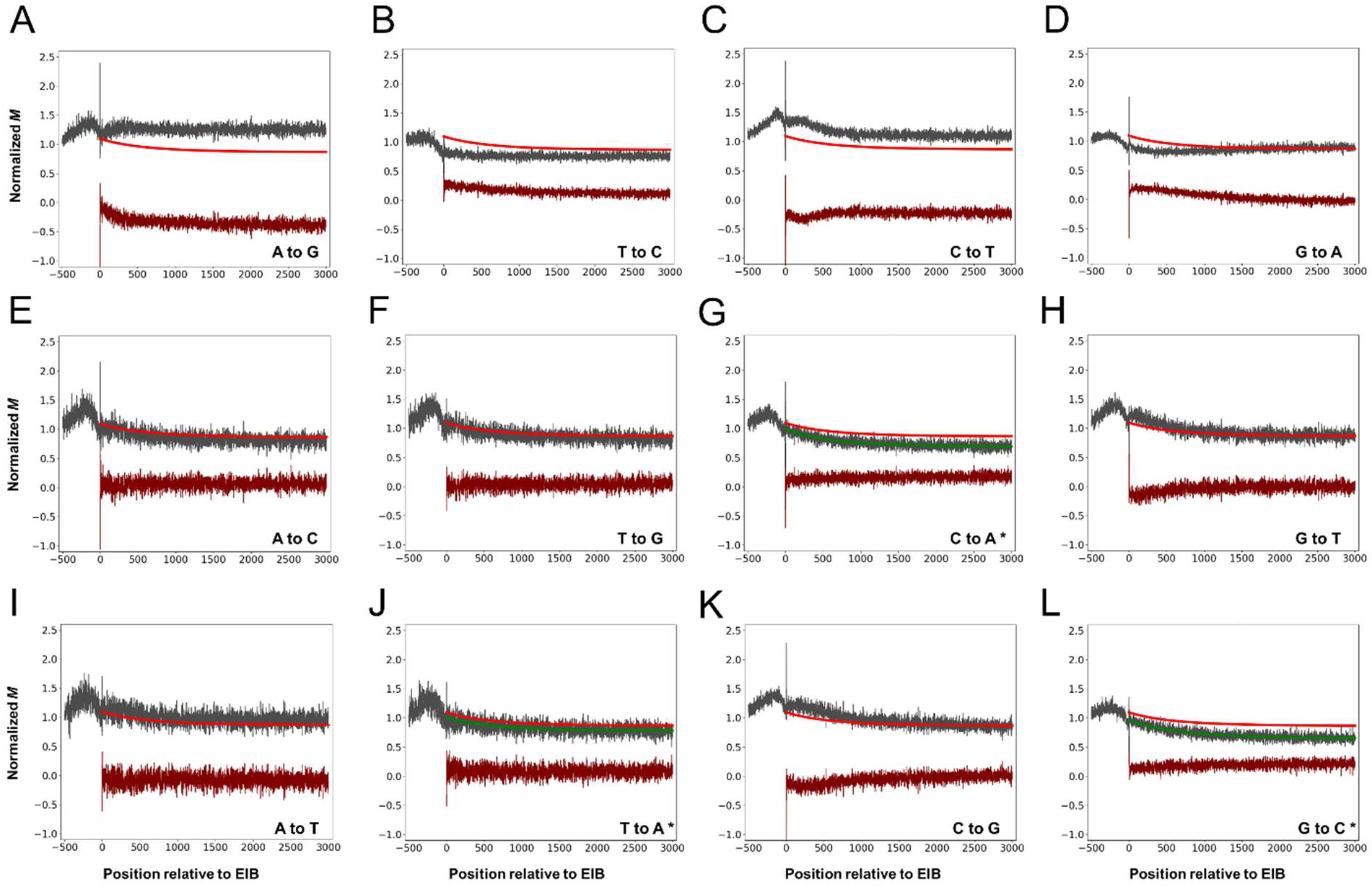
Comparison of overall *M* in first introns with individual rates. A-L) Normalized inferred mutation rates *M* (grey lines) are compared to line fitted overall inferred mutation rate *M* used in the *in silico* time-forward simulation of nucleotide content evolution (smooth bright red line) with the *x-axes* depicting the distance from the exon-intron boundary (EIB). The resulting residual (burgundy line) is also plotted. After AIC analysis, it was determined that incorporating C to A, T to A and G to C residuals significantly improved the simulation performance.

**Supplemental Table 1.**
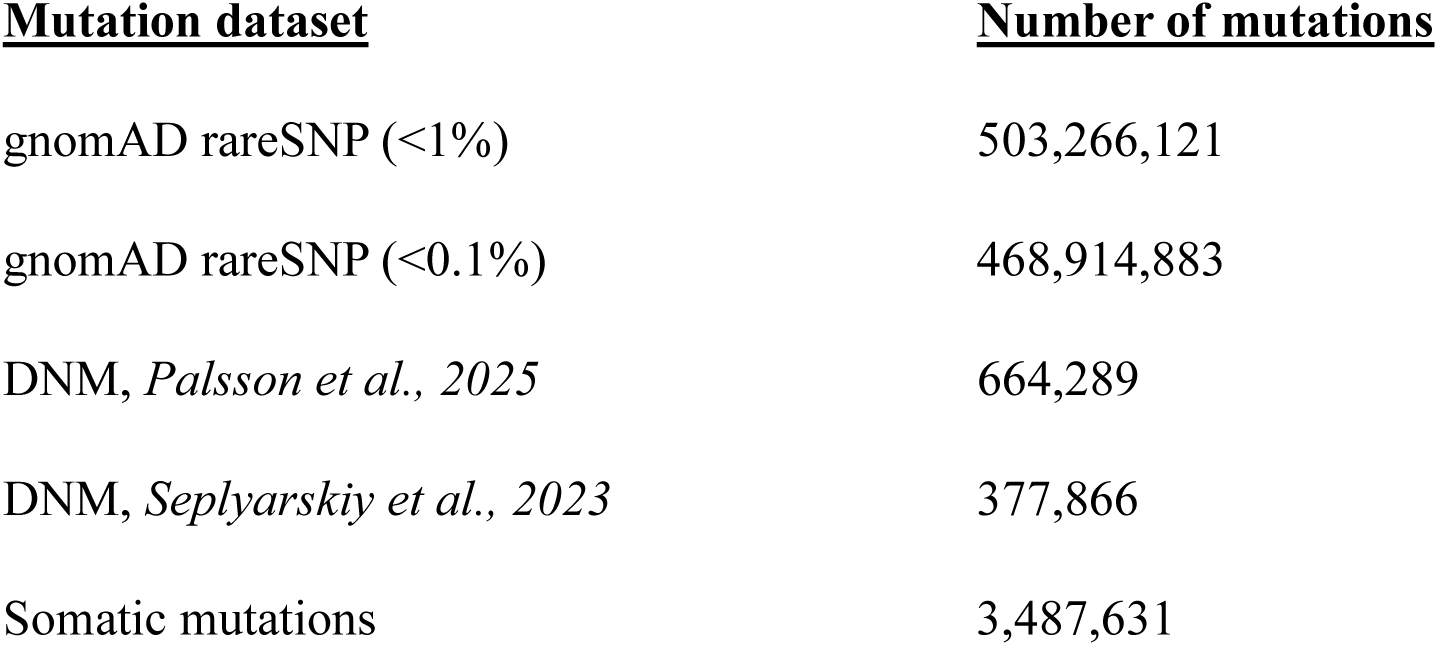
Number of mutations per dataset.

**Supplemental Table 2.** Mispolarization rate of gnomAD reference/alternate alleles across derived allele frequency bins for each analyzed genomic region.

| **DAF bin** | **Intergenic** | **TSS -150**<br>**to +350** | **Exon (TSS**<br>**0 to +350)** | **Intron (TSS**<br>**0 to +350)** | **Recombination**<br>**hotspot** |
| --- | --- | --- | --- | --- | --- |
| **0-0.001** | 0.004% | 0.008% | 0.007% | 0.007% | 0.004% |
| **0.001-0.01** | 0.47% | 0.41% | 0.34% | 0.46% | 0.48% |
| **0.01-0.1** | 3.84% | 3.41% | 3.59% | 3.04% | 3.79% |
| **0.1-0.5** | 24.44% | 23.99% | 23.22% | 23.80% | 25.17% |
| **0.5-0.9** | 66.46% | 66.32% | 66.70% | 65.30% | 67.30% |
| **0.9-1** | 98.98% | 99.10% | 99.35% | 98.73% | 98.68% |

